# A gap-free, telomere-to-telomere genome assembly for the *Caenorhabditis briggsae* reference strain AF16

**DOI:** 10.64898/2026.04.30.721887

**Authors:** Lance M. O’Connor, Nicolas D. Moya, Nikita S. Jhaveri, Robyn E. Tanny, Ayeh Khorshidian, Haimeng Lyu, Helen M. Chamberlin, Scott E. Baird, Erik C. Andersen

## Abstract

The nematode *Caenorhabditis elegans* was the first metazoan to have its genome completely sequenced and assembled. Since that time, researchers have continuously updated the reference genome and manually curated its approximately 20,000 genes. The closely related species, *Caenorhabditis briggsae*, has served as a comparative model because of its similar morphology, mode of reproduction, and patterns of intra-species genetic variation. However, the genomic resources for *C. briggsae* lag behind *C. elegans*, hindering comparative genomics studies between the species. Decades of experimentation have been performed in the AF16 reference strain genetic background, so we obtained high-coverage long-read sequencing and high-throughput chromosome conformation capture data to create an updated reference genome for an isogenic derivative of AF16, named CGC2. The CGC2 genome is vastly improved relative to the existing AF16 assemblies, with no unplaced sequence, no gaps, and telomere-to-telomere contiguity. To provide genomic resources for CGC2, we exploited deep RNA-seq libraries from all developmental stages to predict protein-coding gene annotations comparable in accuracy and completeness to the existing AF16 gene models. We also performed lift-over of 108 validated insertion-deletion variants to the updated coordinate system of the CGC2 genome to facilitate future mappings of mutations. In summary, we present an updated reference genome for the canonical AF16 reference strain with improved genomic resources to enable high-quality intra- and inter-species comparative studies.

## Introduction

Nematodes of the genus *Caenorhabditis* are routinely used to discover genes, genetic interactions, and evolutionarily conserved signaling pathways. *Caenorhabditis elegans*, the most studied species in this genus has pioneered studies of signal transduction, programmed cell death, gene regulation by microRNAs, and gene silencing by RNA interference (Brenner 1974; Ellis and Horvitz 1986; Lee et al. 1993; Wightman et al. 1993; Fire et al. 1998). Furthermore, *C. elegans* became the first metazoan to have its genome fully sequenced (The C. elegans Sequencing Consortium 1998). The research community has since improved the completeness and accuracy of its genome resources (Sönnichsen et al. 2005; Yoshimura et al. 2019; Ichikawa et al. 2025). In recent years, genomic resources and chromosome-level genome assemblies for various *Caenorhabditis* species have been developed (Blaxter et al. 2012; Kanzaki et al. 2018; Stevens et al. 2019; Stevens et al. 2020; Teterina et al. 2020; Noble et al. 2021; Teterina et al. 2023; Kieninger et al. 2024). The availability of many reference genomes enables large-scale comparative studies to elucidate genome evolution, gene-family conservation, and syntenic relationships.

As one of the earlier nematodes used in comparative studies, *Caenorhabditis briggsae* has served as an invaluable model, contributing to our understanding of interspecific differences in development, genomics, and proteomics (Petalcorin et al. 2005; Aevermann and Waters 2008; An et al. 2017; Large et al. 2025). *C. elegans* and *C. briggsae* share morphological traits, ease of cultivation, similar modes of reproduction, amenability to genetic modifications, food preferences, and global distribution (Cutter et al. 2006; Gupta et al. 2007; Félix and Duveau 2012; Schulenburg and Félix 2017; Crombie et al. 2019). However, the molecular pathways underlying similar phenotypes can be different. For example, hermaphroditism evolved independently in *C. elegans* and *C. briggsae*, with divergence at the *tra-2* regulatory level in the sex determination pathway (Nayak et al. 2005; Hill et al. 2006). Additionally, wild strains of *C. briggsae* display higher levels of genetic variation (Cutter et al. 2010; Stevens et al. 2022; Moya et al. 2023) compared to other self-fertilizing species of *Caenorhabditis*, such as *C. elegans* (Lee et al. 2020) or *C. tropicalis* (Noble et al. 2021; Wang et al. 2026).

Despite the importance of *C. briggsae* for comparative analyses, the genomic resources have lagged behind *C. elegans* resources. The reference strain for *C. briggsae*, AF16, was isolated from Ahmedabad, India in 1980 (Fodor et al. 1983). The first draft genome of the AF16 strain was assembled in 2003 by combining whole-genome shotgun sequences and a physical map based on fosmid and bacterial artificial chromosome libraries (Stein et al. 2003). The assembly was updated in 2011 using a genetic map generated from recombinant inbred lines (Ross et al. 2011). The updated genome assembly, “cb4”, is publicly available and routinely used in genomic studies (Ross et al. 2011; Bi et al. 2015; Li et al. 2016; Ren et al. 2018; Ding et al. 2022; Ding et al. 2022; Garrigues and Pasquinelli 2022; Moya et al. 2023; Alkan et al. 2024). However, AF16 cb4 contains many unplaced scaffolds and thousands of gaps (Ren et al. 2018; Stevens et al. 2022; Moya et al. 2023). An improved version of the assembly called “cb5” was released in 2023 (Xie et al. 2024). Although improved, AF16 cb5 still has unresolved gaps and most of the genomic resources, such as gene models are in the AF16 cb4 genomic coordinate system. In contrast to the heavily curated gene annotations for the *C. elegans* reference strain N2, the AF16 cb4 gene annotations are primarily based on homology to N2 and have not been extensively validated. Multiple groups have worked to improve the AF16 gene annotations, but such improvements have remained constrained by the quality of the reference genome (Hillier et al. 2007; Koboldt et al. 2010; Ross et al. 2011; Uyar et al. 2012; Grün et al. 2014; Verster et al. 2014; Bi et al. 2015; An et al. 2017; Ren et al. 2018; Jhaveri et al. 2022).

Our laboratory expanded genomic resources for *C. briggsae* by generating a high-quality chromosome-level genome and manually curated gene models for a new reference strain, QX1410, which is closely related to the AF16 reference strain (Thomas et al. 2015; Stevens et al. 2022; Moya et al. 2023). However, we recognize the need to improve the genome and gene models for the AF16 reference strain because most *C. briggsae* research uses the AF16 strain genetic background (Delattre and Félix 2001; Kirouac and Sternberg 2003; Stein et al. 2003; Cutter et al. 2006; Félix 2007; Hillier et al. 2007; Winter et al. 2007; Cutter and Choi 2010; Woodruff et al. 2010; Félix et al. 2011; Hoyos et al. 2011; Pénigault and Félix 2011; Félix and Duveau 2012; Nuez and Félix 2012; Barkoulas et al. 2016; Félix and Wang 2019; Alkan et al. 2024). Over the last few decades, laboratories have obtained their own copies of the AF16 strain from the *Caenorhabditis* Genetics Center (CGC). Differences in passaging between laboratories have likely led to the accumulation of mutations over time, giving rise to different genotypes of the same strain. Studies in *C. elegans* have shown that the reference strain N2 has different lifespans when collected from different laboratories (Gems and Riddle 2000; Vergara et al. 2009; Sterken et al. 2015). Additionally, isogenic derivatives of the N2 strain exhibit differences in fecundity and developmental time (Mertz et al. 2025). Because of the observed genetic differences among laboratory-specific reference strains, it is important to identify the oldest, historically documented AF16 strain and assemble a reference genome and generate protein-coding gene annotations for this strain.

In this study, we obtained high-coverage PacBio HiFi, Oxford Nanopore Technologies (ONT), and Hi-C sequencing of an isogenic derivative of the AF16 strain, named CGC2, to generate a *de novo* genome assembly. We scaffolded and manually curated the primary CGC2 assembly to produce a gap-free, telomere-to-telomere (T2T) genome. Next, we predicted protein-coding gene models that were comparable in accuracy and completeness to the existing AF16 gene models. We also performed lift-over of 108 validated insertion-deletion (indel) variants (Guo et al. 2009; Guo et al. 2013; Koboldt et al. 2010; Jhaveri and Gupta 2023; Seetharaman et al. 2010; Sharanya et al. 2015; this study) to the updated coordinate system of the CGC2 genome. In summary, we present an updated reference genome for the canonical AF16 reference strain with improved genomic resources to enable high-quality intra- and inter-species comparative studies.

## Materials and methods

### Strain maintenance

PB420 was derived from a precursor of AF16 frozen in 1991. PB420 was thawed in 2020 and had not been grown in long-term culture. It was maintained at 20°C on a modified nematode growth medium (NGMA) (consisting of 1% agar and 0.7% agarose) (Andersen et al. 2014) on 6 cm plates for three generations before collecting samples for RNA extractions. *Escherichia coli* OP50 was used as the food source.

### Short-read Illumina DNA sequencing

DNA was extracted from nematodes grown on three 10 cm NGMA plates when the plates were close to starvation. The animals were washed with M9 into a 15 ml conical tube. The animals were allowed to settle and the supernatant was removed. Fresh M9 buffer was added, and the wash was repeated three times. Genomic DNA was extracted using the Blood and Tissue DNA isolation kit (Qiagen, Catalog #69506) using published protocols (Cook et al. 2016). DNA concentration was determined using the Qubit dsDNA Broad Range Assay kit (Invitrogen, Catalog #Q32853). Sequencing libraries were generated using NEBNext® Ultra™ II FS DNA Library Prep (New England Biolabs, Catalog #E6177L). Samples were sent for sequencing at the Novogene facility, Sacramento, California. All samples were sequenced on the NovaSeq 6000 platform (paired-end 150 bp reads).

### Sample collection for Hi-C, ONT, and PacBio HiFi sequencing

Nematodes were grown on four 10 cm NGMA plates seeded with OP50. When the plates were close to starvation, the animals were washed off with cold (4°C) M9 buffer (3 g of KH2PO4, 6 g of Na2HPO4, and 5 g of NaCl in 1 L Milli-Q water) into a 50 mL conical tube and centrifuged at 2,000 rpm for eight minutes with a break of three for a slow stop of the centrifuge. The supernatant was removed, and the pellet was washed three times using cold M9 buffer + 0.01% Tween 20. A final wash was performed using cold M9 buffer. Most of the supernatant was removed, and the pellet was transferred to a 1.7 mL microfuge tube using a glass pasteur pipette. If smaller pellets were needed, the resuspended material was divided into multiple tubes. The animals were centrifuged at 2,000 rpm for three minutes. Supernatant was removed, and the pellets were weighed. The pellet was flash frozen in liquid nitrogen and stored at −80ºC.

For Hi-C sequencing, 100 mg of the animal pellet was sent to Phase Genomics, Seattle, Washington, where the subsequent steps of crosslinking, extraction, and Illumina short-read paired-end sequencing on the NovaSeq platform were performed. For ONT sequencing, 100 mg of the animal pellet was sent to UC Davis, DNA Technologies Core Facility. DNA was prepared using a standard ONT ligation kit and sequenced using the PromethION platform with a FLO-PRO114M flow cell. For PacBio HiFi sequencing, 60 mg of animal pellet was used to make High Molecular Weight (HMW) DNA using the Monarch HMW DNA Extraction Kit for Tissue (New England Biolabs, Catalog #T3060L). The following modifications were made to the protocol. Before using the pestle, 50 µL of the lysis buffer was added to the sample. The remaining 600 µL was added after homogenization. For the lysis at 56ºC after homogenization, we incubated in a Thermomixer for 15 minutes at 750 rpm and then at 550 rpm for 30 minutes or longer. We used a cold (4°C) Protein Separation Solution. After the inversion step, we left the samples on ice for 10 minutes. We used three Capture beads. For elution, we added 200 µL Elution Buffer, incubated for five minutes at 56ºC at 300 rpm. We then added another 50 µL of the Elution Buffer before separating the DNA from the beads. The final solubilization was done at 37ºC for 30 minutes at 300 rpm. The final samples were sent to Maryland Genomics at the University of Maryland for library generation and sequencing.

### Sample collection for RNA extraction and library prep

RNA samples were collected from PB420 for six stages - embryo, L1, L2, L3, L4, and young adult. Filtration technique was used to obtain the embryos following the established protocol (Jhaveri et al. 2025). For the embryonic stage, embryos were directly harvested after collection by filtration. For the larval stages, the embryos were put on the rotor for 18 - 24 hours, and the L1s were plated on 6 cm NGMA plates the next day. Samples were then collected at specific time points for each larval stage. For the L1 stage, the samples were collected the next day. For the L2, L3, L4, and young adult stages, the samples were collected at L1 + 26.5, L1 + 33.5, L1 + 44.5, and L1 + 49 hours, respectively. The animals were washed twice with M9 buffer to remove the OP50 bacteria and then pelleted by centrifugation at 3,900 rpm for one minute. The supernatant was removed, and Trizol (Ambion, Catalog #15596018) was added at four times the volume of animal pellets to the sample and frozen at −80°C.

### RNA extraction

The staged samples that were frozen in Trizol were thawed at room temperature. RNA was extracted using the chloroform-isopropanol method (Zhang et al. 2022). The RNA pellets were resuspended in nuclease-free water. RNA concentration was determined using the Qubit RNA HS Assay Kit (Invitrogen, Catalog #Q32855), and quality was measured using the Agilent Tapestation 4150. All samples had an RNA integrity number of more than eight and were used to construct RNA-sequencing libraries.

### RNA library preparation for Illumina short read sequencing

The RNA sequencing libraries were prepared in a 12-well strip-tube format. The libraries were prepared from 1 µg of total RNA for each animal stage. mRNA was first purified using the NEBNext Poly(A) mRNA Magnetic Isolation Module (New England Biolabs, Catalog #E7490L). NEBNext Ultra II RNA Library Prep with Sample Purification Beads (New England Biolabs, Catalog #E7775L) were used for RNA fragmentation, first-strand, second-strand cDNA synthesis, and end-repair. The cDNA libraries generated were then ligated with adapters and unique dual indexes from the NEBNext Multiplex Oligos for Illumina (New England Biolabs, Catalog #E6443A) and amplified using 12 PCR cycles. The manufacturer’s protocols were followed for all steps. The concentration of each library was measured using the Qubit dsDNA BR Assay Kit (Invitrogen, Catalog #Q32853). All six libraries were pooled and run on a 2% agarose gel (170 V) for 40 to 45 minutes. Gel extraction and purification of the required band size were carried out using the QIAquick gel extraction kit (Qiagen, Catalog #28706). Concentration of the purified sample was determined using the Qubit dsDNA BR Assay Kit (Invitrogen, Catalog #Q32853). The sample (4 nM, total volume of 40 µL) was sent to Johns Hopkins Genetic Resources Core Facility (GRCF), Baltimore, Maryland, for sequencing. A total of 12 short-read RNA-sequencing libraries (two for each stage) were sequenced.

### Comparison of AF16 derivatives

Our lab has a collection of eight AF16 derivative strains obtained from different laboratories (Supplementary Table 2). We aligned the genomes of these AF16 derivatives to the QX1410 reference genome using the Andersen laboratory Nextflow alignment pipeline (https://github.com/AndersenLab/alignment-nf). To compare these AF16 derivatives, we performed variant calling on the sequencing alignment (BAM) files using the Andersen laboratory Nextflow GATK pipeline (https://github.com/AndersenLab/wi-gatk). The run generated a joint variant call file (VCF) containing all homozygous bi-allelic SNVs across all AF16 derivatives. The number and proportion of identical SNV matches between all strain pairs was generated with *gtcheck* from bcftools vX (Danecek et al. 2021) (custom parameters: *-u GT,GT*). A genotype matrix of all SNVs across the eight AF16 derivative strains was extracted using a custom bash script. We queried the genotype matrix to identify the number of private and shared alleles at SNV sites with perfect information (no missing genotypes) across the AF16 derivatives using a custom R script. Custom scripts are available on GitHub (see Data Availability).

### Genome assembly

Sequenced PacBio HiFi genomic reads were demultiplexed and adapters were removed by the University of Maryland sequencing core using *lima* v2.12.0 (https://lima.how). PCR duplicates were removed using *pbmarkdup* v1.1.1 (https://github.com/PacificBiosciences/pbmarkdup). A primary *de novo* assembly was generated from the deduplicated HiFi reads and ONT sequencing reads that passed quality control (a mean Phred score > 10) using *hifiasm* v0.24.0-r702 (Cheng et al. 2021) with the initial bloom filter *-f39*, which is recommended to save memory on k-mer counting for large amounts of sequencing data, haplotig purging disabled (*-l0*, recommended for homozygous genomes), and ONT reads provided to the *-ul* argument. After assembly, contigs were taxonomically classified using *DIAMOND* 2.1.13 (Buchfink et al. 2021) with parameters *--faster* and an e-value of 1e-10 against the UniProt database (Release 2025_03) (UniProt Consortium 2025). Contigs that were annotated as non-*Nematoda* were purged using *BlobToolKit* v4.4.5 (Challis et al. 2020) with *--param bestsumorder_phylum--Inv=Nematoda*, producing a final assembly consisting of 13 contigs (Supplementary Figure 3). *pbmarkdup, hifiasm*, and *BlobToolKit* executions were carried out using the Nextflow pipeline available at https://github.com/AndersenLab/assembly-nf.

### Genome scaffolding and quality control

Alignment of Hi-C data to the contiguous assembly produced by *hifiasm* was performed using *bwa-mem2* v2.2.1 (Vasimuddin et al. 2019) with parameters *-5SP*, and *Picard* v3.4 (https://broadinstitute.github.io/picard/). MarkDuplicates was used to mark duplicate Hi-C reads. Scaffolding of the contiguous assembly produced by *hifiasm* using Hi-C reads was performed using *YaHS* v1.2.2 (Zhou et al. 2023) with default parameters. Seven scaffolds were produced, with a total of six gaps and zero contig breaks. Six of the seven scaffolds were assigned to QX1410 chromosome IDs based on primary alignment to QX1410 chromosomes using *NUCmer* v3.1 (Marçais et al. 2018). The remaining unplaced scaffold, which consisted of a single contig, scaffold 7 (58 kb), was deemed a duplicate haplotig and removed from the scaffolded assembly. This decision was based on scaffold 7’s primary alignment to a different scaffold (scaffold 4), HiFi reads multimapping to both the scaffold 4 and scaffold 7, and approximately half the Hi-C coverage compared to other scaffolds (Supplementary Figure 4).

### Chromosome gap closing

To close the remaining gaps in the scaffolded assembly, we manually inspected HiFi and ONT read alignments produced using *minimap2* v2.28-r1209 (Li 2018) at the gap loci in the Integrative Genomics Viewer (IGV) v2.19.7 (Robinson et al. 2011). Five gaps were located on the left-end of chromosome V in an rDNA repeat sequence, previously characterized in QX1410 (Stevens et al. 2022), and one gap was located on chromosome III at position 10,743,295 to 10,743,394 (Supplementary Table 4). The gap on chromosome III was closed manually based on evidence from HiFi and ONT read alignments that supported the deletion of the YaHS-inserted Ns and 10 bp upstream (Supplementary Figure 5). To manually assess the gaps on the left-end of chromosome V, a Hi-C contact map was created using *Juicer* v1.22.01 (Durand et al. 2016) and visualized with Juicebox (development version v2.15) (Durand et al. 2016). Telomeric repeat sequence was identified in an internally placed contig and was manually placed at the end of chromosome V (Supplementary Figure 6). *TGS-GapCloser* v1.2.1 (Xu et al. 2020) was then used to close the remaining five gaps on the left-end of chromosome V with ONT reads and three rounds of polishing performed using *Pilon* v1.24 (Walker et al. 2014) using short-read sequencing data (Supplementary Figure 7). We did not re-orient rDNA contigs that had inverted alignments to the QX1410 rDNA repeat sequence because we did not observe substantial evidence for their orientation based on ONT read alignment or Hi-C contacts. The final chromosomal scaffolds after gap closing were reoriented to match QX1410 chromosome orientations using *seqkit* seq v2.10.0 (Shen et al. 2016) with parameters *-rp*. Genome completeness was assessed using genome BUSCO v5.8.3 (Simão et al. 2015) with the *nematoda_odb10* database that contains 3,131 single-copy orthologs found in the *Nematoda* phylum. Statistics of QV and sequence completeness were calculated using *Merqury* v1.3 (Rhie et al. 2020) with Illumina short-reads and 21-mers created using *Meryl* v1.4.1 (Rhie et al. 2020) (Supplementary Table 6). Telomeric repeat sequence was detected using *seqkit* locate v2.10.0 (Shen et al. 2016) and repeat count was quantified per kb across the CGC2 genome using *bedtools* v2.31.0 (Quinlan and Hall 2010) (Supplementary Figure 11). Final genome statistics were calculated using *bbtools* v37.62 (Bushnell 2014) *stats*.*sh* script with parameters *–format=6* and *–gcformat=0*. The mitogenome assembly of AF16 available from NCBI accession NC_009885 was concatenated in the final nuclear genome assembly.

### Insertion-deletion variant position liftover

We experimentally validated 17 indel markers in the AF16 cb4 genome. We also consolidated 91 indels that had been experimentally validated by other groups (Supplementary Table 7) in the AF16 cb4 genome. We created a FASTA file containing the forward and reverse primer sequences of each indel site. We used *blastn* from BLAST v2.16.0+ (Camacho et al. 2009) to identify the positions of significant primer hits to both the AF16 and CGC2 genomes (custom parameters: *-task blastn-short -dust no -soft_masking false -outfmt 6*). We excluded any hits with more than one nucleotide mismatch. Processing of BLAST results was carried out using a custom R script available on GitHub (see Data Availability). Indel positions were directly lifted over from the genomic positions of mapped primer pairs when these primer pairs had unique sequence matches to both genomes (92 primer pairs with perfect sequence matches in both genomes; three primer pairs had one mismatch in one primer in both genomes). One additional indel (cb-m179) was also directly lifted over from the genomic positions of its mapped primer sequences to the CGC2 genome despite being mapped to two possible positions in the AF16 cb4 genome. For the remaining 12 indels, we performed the lift over for nine primer pairs in which no more than one primer aligned to multiple genomic positions in both genomes, assigning the position of that primer with multiple genomic positions based on its proximity to its paired primer with a unique genomic position. The three remaining primer pairs mapped to multiple genomic positions in both genomes and could not be lifted over.

### Protein-coding gene annotation

Protein-coding gene annotation for CGC2 was generated using *BRAKER3* v3.0.8 (Gabriel et al. 2023) and the 12 stage-specific RNA libraries described previously along with a custom library of protein sequences from *C. elegans* N2 (PRJNA13758) and *C. briggsae* QX1410 (PRJNA784955) as evidence. RNA reads were aligned to the CGC2 genome using *STAR* v2.7.11b (Dobin et al. 2013) in basic two-pass mode with custom parameters for compatibility with *BRAKER3* (*--outSAMstrandField intronMotif*) and a maximum intron size of 10 kilobases (*--alignIntronMax 10000*). Repetitive sequences were masked in CGC2 genome using *RepeatMasker* v4.2.1 (Smit et al. 2015) and a custom *C. briggsae* repeat library described in a previous study (Moya et al. 2025). Gene model biological completeness of each set of gene models was estimated using protein BUSCO v5.8.3 (Simão et al. 2015) with the *nematoda_odb10* database. *RepeatMasker, BRAKER3*, and BUSCO executions were carried out using a Nextflow pipeline available at https://github.com/AndersenLab/geneAnno-nf on branch “CGC2_geneAnno”. Mitochondrial genes were retrieved from NCBI accession NC_009885.

### Analysis of gene annotation completeness and accuracy

We identified putative orthologous groups using *OrthoFinder* v3.1.0 (Emms et al. 2025) with protein sequences translated from gene annotations of AF16 (PRJNA10731), QX1410 (PRJNA784955, (Moya et al. 2023), CGC2 (this study), and N2 (PRJNA13758). Only the longest isoform of each gene was used, selected with *agat_sp_keep_longest_isoform*.*pl* from *AGAT* v1.5.1 (https://zenodo.org/records/16317950). A total of 96,134 protein sequences across all four gene annotations were clustered into 19,519 orthologous groups. We queried the resulting orthologous groups table to record the number of single-copy orthologs shared across at least two *C. briggsae* gene annotations (absent or present as a single copy in the third *C. briggsae* annotation). We used *Liftoff* v1.6.3 (Shumate and Salzberg 2021) to recover the count and positions of gene models absent in one *C. briggsae* annotation but present in others. For single-copy orthologs present across AF16, CGC2, and N2 gene annotations (8,021 genes, Supplementary Table 12), we estimated percent protein length difference from N2 by the ratio of a *C. briggsae* (CGC2 or AF16) protein sequence length and the protein sequence length of its N2 ortholog minus one. For each *C. briggsae* annotation, we classified a protein sequence as “identical” if the estimated percent protein length difference from N2 was equal to 0%, as “near-identical” if between −5% and 5% (inclusive) difference, or as “deviated” if less than −5% or greater than 5% difference. We classified deviated protein sequences as “concordant” if lengths between the AF16 and CGC2 gene annotations were within 5% of each other or otherwise as “discordant”. Processing of orthologous groups and protein sequence length comparisons and classifications were carried out using a custom R script available on GitHub (see Data Availability).

## Results

### Genome quality issues in the current AF16 reference genome

Our laboratory previously published a high-quality reference genome and gene models for QX1410, a wild strain related to AF16 (Stevens et al. 2022; Moya et al. 2023). The QX1410 reference genome revealed the pervasive issues associated with the AF16 cb4 genome assembly. Previous studies reported that the AF16 cb4 assembly contains miscaffolded regions (Ren et al. 2018; Stevens et al. 2022; Moya et al. 2023), which cause many artifactual inversions in AF16 relative to QX1410 and lead to inappropriate conclusions about synteny and gene order (Fig. 1a). Of the 47 inverted alignments in AF16 relative to QX1410, 31 are larger than 100 kb, with an average size of 167 kb (Stevens et al. 2022) (Fig. 1a, Supplementary Table 1). By contrast, the genome assembly of a genetically divergent wild strain, VX34, has only one inverted alignment compared to QX1410 that is greater than 100 kb (Stevens et al. 2022). Although some of these inverted alignments in AF16 might be biologically accurate, the inflated number is likely because researchers scaffolded AF16 from a highly fragmented assembly (5,074 contigs, 47 kb N50, Table 1). The genome assembly errors of AF16 extend further, with 4,707 gaps and 361 unplaced scaffolds (3,201,015 bps of sequence, Fig. 1b), in comparison to QX1410 which has seven gaps and no unplaced sequence. The genome assembly of AF16 has a BUSCO score of 98.8%, which indicates that a small proportion of single-copy orthologs conserved in the *Nematoda* phylum are missing. By proxy, the missing BUSCOs likely reflect missing genes across the entire gene annotation of AF16 caused by the joint effects of gene prediction error and genome assembly quality, as described in previous observations made using comparative analyses of gene annotations in *C. briggsae* (Moya et al. 2023; Lyu et al. 2024). The presence of thousands of gaps, over three Mb of unplaced sequence and misoriented sequences in the AF16 cb4 genome makes comparative genomic studies challenging, and can lead to incorrect measurements of synteny, structural variation, and absent gene content. Since the release of the AF16 cb4 genome assembly in 2011, researchers have worked to improve this assembly (Ren et al. 2018; Xie et al. 2024), with the release of the AF16 cb5 genome in 2023. Although the AF16 cb5 genome assembly is drastically more contiguous than AF16 cb4, it still contains 24 gaps, an inversion greater than 495 kb on the right-end of chromosome IV relative to QX1410, and lacks nearly all telomeric chromosome ends (Table 1, Supplementary Figure 2 and 10).

**Fig. 1.**
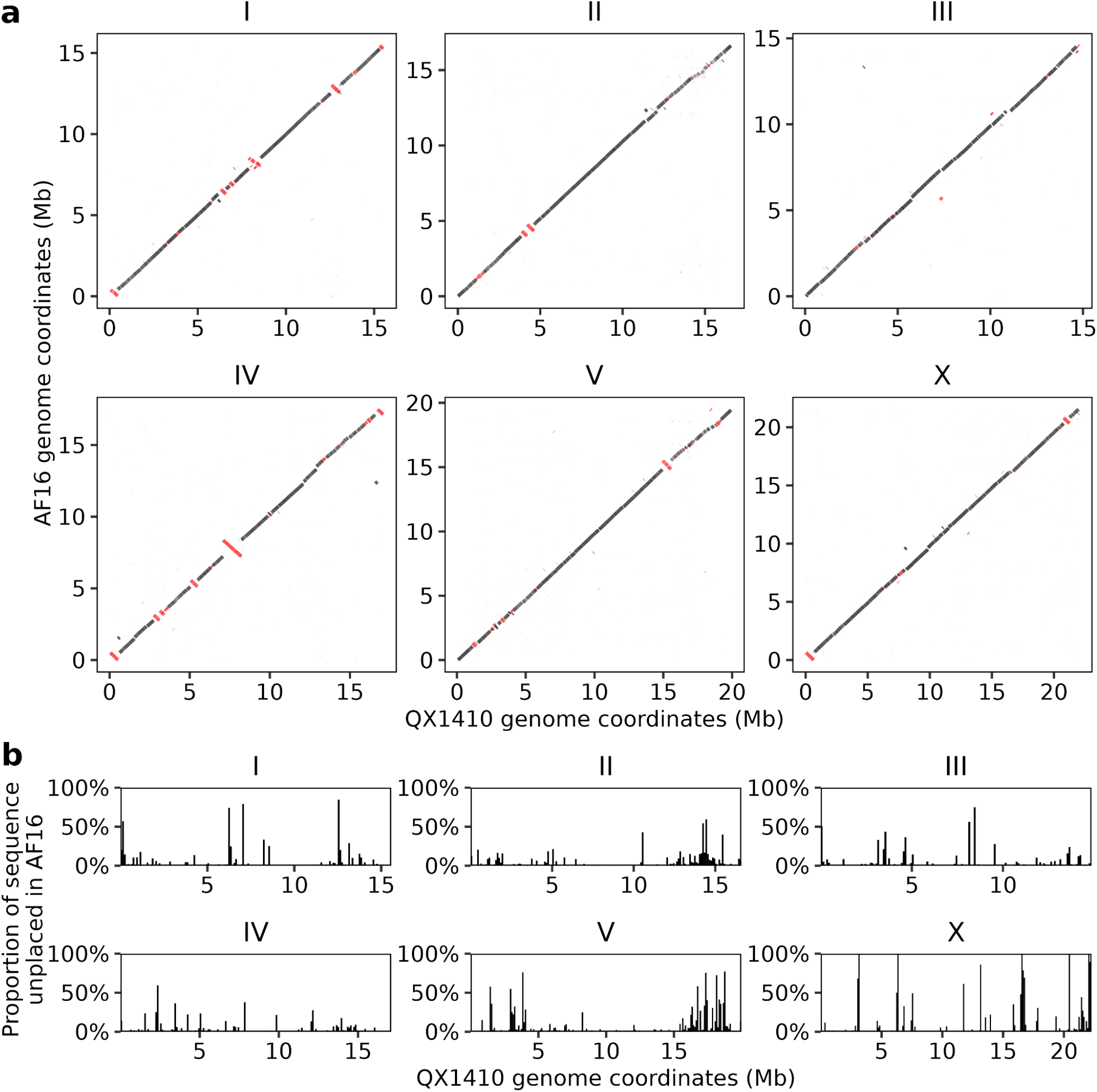
Whole-genome alignment of AF16 cb4 to QX1410. (a) Colinearity of AF16 cb4 to QX1410 genomes generated using nucmer. Inter-chromosomal alignments and alignments spanning less than 1 kb are not shown. Alignments in the reverse orientation are displayed in red. (b) The proportion of sequence that is contained in AF16 cb4 unplaced scaffolds per 100 kb window across the QX1410 genome.

**Table 1.** Genome assembly statistics for *C. briggsae* reference genomes.

|  | **AF16 - cb4\*** | **AF16 - cb5** | **CGC2** | **QX1410\*** |
| --- | --- | --- | --- | --- |
| **Accession** | PRJNA10731 | PRJNA917437 | PRJNA1451919 | PRJNA78495 |
| **Version** | WS276 | v1 | v1 | v1 |
| **Size (Mb)** | 108.4 | 105.5 | 106.8 | 106.2 |
| **Number of Scaffolds** | 367 | 6 | 6 (+MT) | 6 (+MT) |
| **Number of unassigned scaffolds** | 361 | 0 | 0 | 0 |
| **Percentage of assembly span in six scaffold (%)** | 97.0 | 100 | 100 | 100 |
| **Scaffold N50 (Mb)** | 17.5 | 17.22 | 17.39 | 17.1 |
| **Number of contigs** | 5,074 | 30 | 12 | 14 |
| **Contig N50 (Mb)** | 0.05 | 6.18 | 17.39 | 14.7 |
| **Number of gaps** | 4,707 | 24 | 0 | 7 |
| **Span of Ns (kb)** | 2,965.5 | 5.2** | 0 | 3.5 |
| **BUSCO completeness (%)** | 98.8 | 99.7 | 99.7 | 99.4 |
| **QV score** | - | 46.45 | 54.47 | 45.6 |

\*Data from Stevens *et al.*, 2022.
\*\*24 unique spans of N ranging from 10 - 500 bases.

Although the chromosome-level assembly and manually curated gene annotations for QX1410 provide a valuable resource for comparative genomic studies, the genetic background of the *C. briggsae* reference strain, AF16, has historically been used in laboratory experiments. Since the first isolation of the AF16 strain in 1980 (Fodor et al. 1983), the CGC cryopreserved and disseminated it to different laboratories all around the world. The culture, maintenance, and passage conditions and protocols of laboratories working with AF16 might differ, which give rise to genetic differences among AF16 strains that accumulate over time. Among the eight AF16 isogenic derivative strains that we requested from different laboratories (Supplementary Table 2), we selected the PB420 strain as the new AF16 reference because of its documented historical provenance. An analysis of genome sequences across all AF16 derivative strains later revealed that the PB420 strain only carries 24 private alleles relative to all other AF16 derivative strains, nearly a 10-fold reduction relative to the AF16 derivative strain (224 private and potential laboratory derived alleles) that we obtained from CGC in 2009 (see Methods, Supplementary Figure 1, Supplementary Table 3). A few notable AF16 derivative strains have fewer counts of private alleles when compared to PB420 (Supplementary Table 3), but these strains showed narrow differences in private allele counts and oftentimes had less clear provenance. We will hereafter refer to the new AF16 reference strain, PB420, as CGC2. This strain is available for future studies from the *Caenorhabditis* Genetics Center.

### CGC2, an updated AF16 *C. briggsae* reference genome

To create a new reference genome for AF16, we generated high-coverage long-read sequencing data for CGC2 using PacBio HiFi (~61x coverage) and ONT using the PromethION platform (~1,048x coverage) and sequenced chromosome-conformation capture (Hi-C) libraries using Illumina technology (~398x coverage). We removed HiFi read duplicates and performed joint *de novo* assembly of HiFi and ONT sequence data using hifiasm (Cheng et al. 2021), producing 35 contigs (Supplementary Figure 3a). Twenty of the 35 contigs were small (approximately 2 Mb total), contained high GC content, and were classified as *Pseudomonadota* by DIAMOND (Buchfink et al. 2021). Another two contigs were classified as *Ascomycota* (approximately 145 kb total). These 22 contigs were removed from the primary assembly using BlobToolKit (Challis et al. 2020) (Supplementary Figure 3b), which produced a contiguous assembly of 13 contigs (17.39 N50, Table 1). These 13 contigs were then scaffolded into chromosomes using Hi-C data, which generated seven scaffolds with no contigs being broken. Six of the seven scaffolds represented nuclear chromosomes, with a single scaffold identified to be a duplicate haplotig (see Methods, Supplementary Figure 4), which we purged from the assembly. After scaffolding, six gaps remained, with five of the six gaps in the first 350 kb of sequence on the left-end of chromosome V, and one gap at approximately 10.74 Mb on chromosome III (Supplementary Table 4). The gap on chromosome III was manually closed by assessing short-read, HiFi, and ONT alignments that spanned the gap (Supplementary Figure 5). To assess the gaps on the left-end of chromosome V, we created a Hi-C contact map and found an internal contig that contained telomeric repeat sequence. This contig was manually re-positioned to the left-end of chromosome V (Supplementary Figure 6-7) based on Hi-C contact support. The remaining five gaps on the left-end of chromosome V were closed using TGS-GapCloser (Xu et al. 2020) (see Methods, Supplementary Figure 7).

The final CGC2 genome is 106.8 Mb in total span, with similar chromosome sizes to the existing AF16 cb4 reference genome (Table 1, Supplementary Figure 8) and has no unplaced sequences contained in six chromosome scaffolds. CGC2 is much more contiguous than the AF16 cb4 assembly (12 contigs, N50 of 17.39 Mb compared to 5,074 contigs, N50 47 kb (Table 1)). The BUSCO completeness score of CGC2 is 99.7% (with 0.3% duplication), an improvement from the previous score of 98.8% in AF16 cb4, indicative that CGC2 has captured nearly all single-copy orthologs conserved in the *Nematoda* phylum (Table 1). Similar to the divergent strain VX34, CGC2 only has one inversion larger than 100 kb in relation to QX1410. This inversion is on the right arm of chromosome V in contiguous CGC2 sequence, which indicates that it is not an artifact of a misoriented contig from scaffolding (Fig. 2). Furthermore, this inversion is at the same genomic locus in the VX34, AF16 cb4, AF16 cb5, and CGC2 genomes (Supplementary Figure 9). The genomic region in QX1410 of this inversion does not contain any gaps and 1,255 bps of sequence flanking the inversion breakpoints are reverse-complements of each other (Supplementary Figure 10a). Additionally, QX1410 ONT read alignments against the QX1410 genome assembly span the inversion breakpoints loci, and CGC2 HiFi read alignments at the inversion breakpoints are hard-clipped, which supports that the right-arm of chromosome V is correctly assembled and indicates that QX1410 most likely has a private inversion allele at this locus (Supplementary Figure 10b). Lastly, a QX1410 genetic map constructed using recombinant inbred lines between VX34 and QX1410 displayed no detectable recombination in this approximately 625 kb inversion (Stevens et al. 2022).

**Fig. 2.**
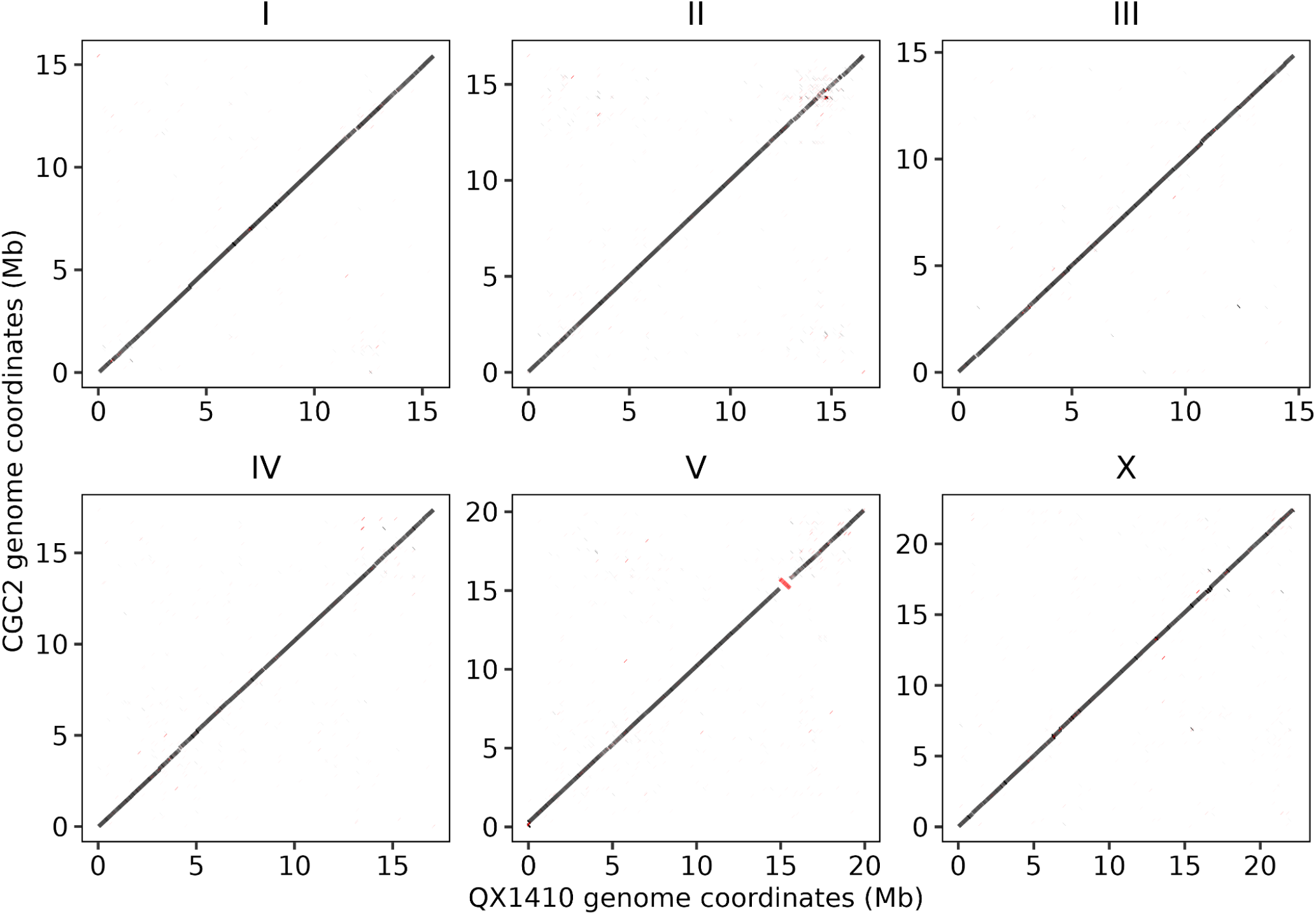
Whole-genome alignment of CGC2 to QX1410. Colinearity of CGC2 to QX1410 genomes generated using nucmer. Inter-chromosomal alignments and alignments spanning less than 1 kb are not shown. Alignments in the reverse orientation are displayed in red.

To further assess the quality of the CGC2 genome, we used Merqury (Rhie et al. 2020) and short-read sequencing data to estimate a sequence completeness of 98.97% and a genome-wide QV score of 54.47, which corresponds to approximately one base error every 280,000 bases (Table 1, Supplementary Table 6). We validated the telomeric repeat sequence ([TTAAGGC]*n*) at the ends of every chromosome in CGC2, which are absent in both of the previous AF16 assemblies and are absent or collapsed in three chromosomes in the QX1410 genome assembly (Supplementary Figure 11). To assess the five gaps closed on the left-end of chromosome V, we aligned HiFi and ONT sequencing data and found the average HiFi fold-coverage in the first 335 kb of sequence is approximately 252x, compared to the chromosome V average of approximately 60.1x, which indicates a potentially collapsed region (Supplementary Figure 12). This region contains an rDNA cluster that previous studies characterized in *C. elegans*, which contains ~55 repetitive cistron units that sit adjacent to the telomeric repeat sequence on chromosome I (Ellis et al. 1986; The C. elegans Sequencing Consortium 1998). In QX1410, this rDNA cluster is also collapsed on the left-end of chromosome V with no telomeric repeat sequence (Supplementary Figures 7 and 11) (Stevens et al. 2022). After completion of the CGC2 genome, whole-genome alignments of AF16 cb4 to CGC2 have similar colinearity to AF16 cb4 to QX1410 (Supplementary Figure 13, Fig. 1). However, the QX1410 private inversion allele on the right-arm of chromosome V is not present in AF16 cb4 in relation to CGC2.

### Insertion-deletion position liftovers for the CGC2 strain

A polymorphic map was previously constructed using the AF16 reference strain and a wild strain with a divergent genetic background, HK104 (Koboldt et al. 2010). The map encompasses experimentally validated single nucleotide variants and indels (7-2,000 bps) that have been instrumental in the identification of mutations underlying mutant phenotypes (Guo et al. 2009; Seetharaman et al. 2010; Guo et al. 2013; Sharanya et al. 2015; Jhaveri and Gupta 2023). We lifted over the genomic positions of 108 validated indels (Guo et al. 2009; Koboldt et al. 2010; Seetharaman et al. 2010; Guo et al. 2013; Sharanya et al. 2015; Jhaveri and Gupta 2023) from AF16 cb4 to the newly assembled CCG2 genome using BLAST (Camacho et al. 2009) to map the genomic positions of indel primer sequences to both the AF16 cb4 and CGC2 genomes (Fig. 3, Supplementary Table 7). Allowing up to one nucleotide mismatch, we identified mappable positions for all 108 indel primer pairs in both the AF16 cb4 and CGC2 genomes. From these 108 primer pairs, 92 mapped to unique genomic positions with no mismatches to both genomes (Supplementary Table 8). Additionally, three primer pairs mapped to unique genomic positions but exhibited a single-nucleotide mismatch in a single primer in both AF16 cb4 and CGC2 (Supplementary Table 8). One additional primer pair (cb-m179) mapped to unique positions only in the CGC2 genome (Supplementary Table 8). Among the remaining 12 primer pairs, nine pairs included one primer with a single mapped genomic position and the other primer with multiple possible mapped genomic positions in both genomes (Supplementary Figure 14). We inferred the most likely position of the multi-mapping primer based on its proximity to the single-mapping primer (Supplementary Table 8). The last three primer pairs could not be lifted over because both primer sequences mapped to multiple genomic positions in both genomes (Supplementary Figure 14, Supplementary Table 9). The updated positions of the markers in the CGC2 genome will be useful in genetic mappings to approximate the positions of mutations and detect recombination events between CGC2 and other strains where indel genotypes differ.

**Fig. 3.**
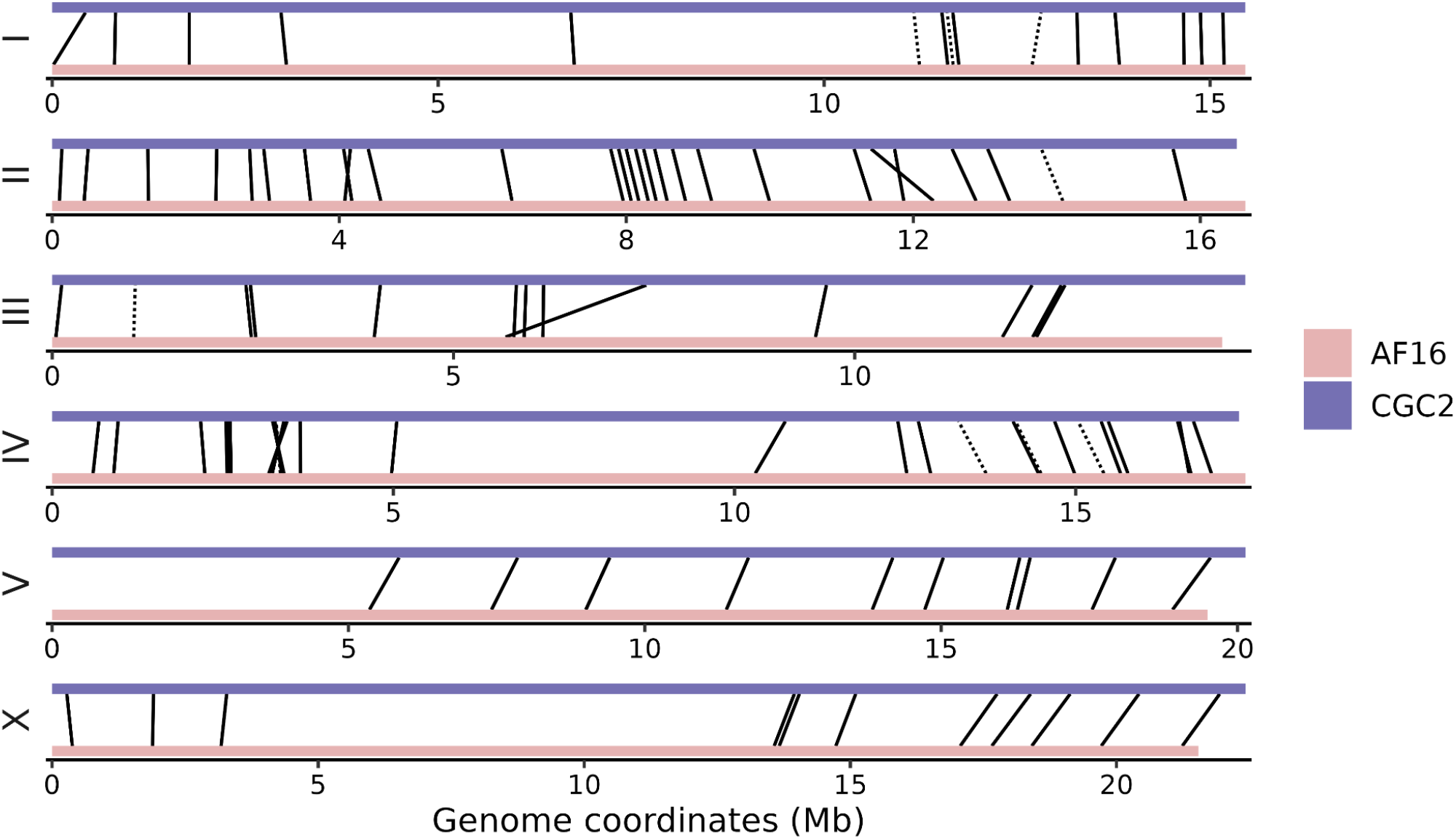
Position liftovers of validated insertion-deletion variants from AF16 cb4 to CGC2. Each rectangle represents a *C. briggsae* chromosome colored by reference genome and organized by chromosome ID across the y-axis. Physical position in mega bases is displayed on the x-axis. Solid lines represent the positions of primer pairs that map uniquely to both AF16 cb4 and CGC2 genomes. Dotted lines represent primer pairs where the position of one primer was inferred from multiple mapping positions in both genomes based on proximity to its paired primer, which mapped to a unique genomic position.

### Protein-coding gene annotation for the CGC2 strain

To produce a high-quality protein-coding gene annotation for the CGC2 genome, we first sequenced its transcriptome in twelve developmental stage-specific libraries (two libraries for each developmental stage, see Methods). We estimated an average fold-coverage of approximately 63x per library (Supplementary Table 10) using only cDNA reads that mapped to unique genomic locations and an estimated exome size of 35 Mb. We leveraged cDNA alignments and a custom library of *C. elegans* (N2 strain) and *C. briggsae* (QX1410 strain) protein sequences as evidence to predict protein-coding genes in the CGC2 genome using BRAKER3 (Gabriel et al. 2023) (see Methods). The BUSCO completeness of these protein-coding gene annotations was 99.4%. We further investigated the biological completeness of these protein-coding gene annotations by identifying and comparing single-copy orthologs relative to the existing AF16 cb4 gene annotations (PRJNA10731), the manually curated genes for the QX1410 strain (Moya et al. 2023) (PRJNA784955), and the *C. elegans* N2 gene annotations (PRJNA13758). We queried a total of 19,519 putative orthologous groups generated using OrthoFinder (Emms et al. 2025) and identified a total of 15,956 single-copy orthologs between any two *C. briggsae* gene annotations. A small proportion of these single-copy orthologs (183/15,956) map between different chromosomes across *C. briggsae* genomes (Supplementary Figure 15). These inter-chromosomal mappings of single-copy orthologs appear in excess relative to AF16 cb4 (Supplementary Figure 15), consistent with misplaced contigs in the AF16 cb4 genome. Of all 15,956 single-copy orthologs, 89.4% (14,258) were present across all three *C. briggsae* gene annotations (Fig. 4a, Supplementary Table 11) and 10.6% (1,698) were absent in one gene annotation (Fig. 4b-c, Supplementary Table 11). The majority of these absences (56.7%, 964/1,698) were missing in the AF16 cb4 gene annotations (Fig 4c), consistent with previous observations of incomplete gene content associated with assembly errors in the current AF16 cb4 genome (Stevens et al. 2022; Moya et al. 2023; Lyu et al. 2024). Moreover, we were unable to identify *C. elegans* orthologs to the vast majority of the recorded gene absences in AF16 cb4 (90.1%, 869/964) that are present in CGC2 and QX1410 (Supplementary Table 11), suggesting that novel genes in CGC2 represent genes specific to *C. briggsae*. Of the remaining recorded absences, 24.4% (414/1698) were missing in the CGC2 gene annotations and 18.8% were missing in the QX1410 gene annotations. In contrast to AF16 cb4 absences, we failed to identify N2 orthologs for a smaller proportion of absences (61.8%, 282/414) associated with the CGC2 gene annotations, perhaps reflecting a stronger adherence to *C. elegans* homology data in the AF16 cb4 gene annotations. We used Liftoff (Shumate and Salzberg 2021) to test if the novel genes in CGC2 could be found in the AF16 cb4 genome, where we were able to recover genomic positions for 99% (860/869) of absent genes in AF16 cb4 that lack an N2 ortholog. Similarly, we were able to recover 83% (234/282) of absent genes in CGC2 that lack an N2 ortholog and are present in AF16 cb4 (Supplementary Table 11). These results indicate that both AF16 cb4 and CGC2 gene annotations have comparable sensitivity to predict genes shared with the N2 strain, but contain different sets of unique genes that likely reflect differences in gene prediction methodology and evidence, where AF16 cb4 displays a more limited sensitivity of *C. briggsae* specific gene discovery than the CGC2 gene annotations.

**Fig. 4.**
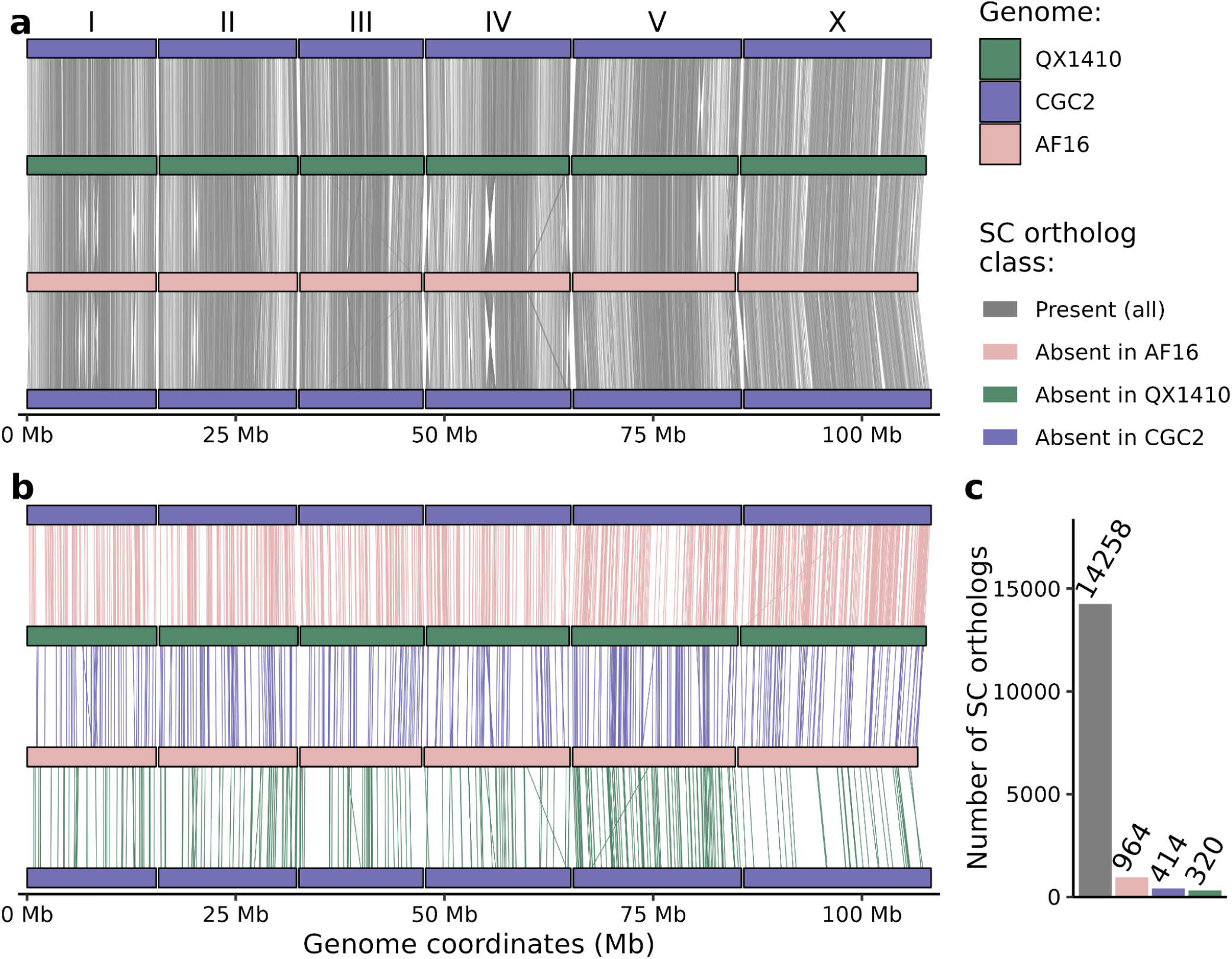
Single-copy (SC) ortholog mapping between *C. briggsae* reference genomes. (a,b) Each rectangle represents a *C. briggsae* chromosome colored and organized in rows by reference genome (CGC2, purple; QX1410, green; AF16 cb4, pink). The CGC2 genome is shown on the top and bottom rows to display all pairwise genome mappings. Physical positions in megabases (Mb) are displayed on the x-axis. Lines between rectangles indicate the positions of intra-chromosomal single-copy ortholog mappings between two genomes. (a) SC orthologs present in all three *C. briggsae* genomes are shown as lines colored in grey. (b) SC orthologs present in CGC2 and QX1410 but absent in AF16 cb4 are shown as lines colored in pink, SC orthologs present in CGC2 and AF16 cb4 but absent in QX1410 are shown in green, and SC orthologs present in AF16 and QX1410 but absent in CGC2 are shown in purple. (c) Bar chart showing counts of SC orthologs present in all *C. briggsae* gene annotations or absent in one annotation. Bars showing counts of absences are colored by the reference genome. These counts include intra-chromosomal mappings in panels (a) and (b) as well as a few inter-chromosomal mappings (Supplementary Figure 15).

We also assessed the accuracy of the CGC2 gene annotations by performing a comparison of protein sequence lengths from conserved single-copy orthologs between *C. briggsae* (CGC2 and AF16 cb4) and *C. elegans* (N2). Using the N2 protein sequences as a benchmark dataset, deviations in protein sequence length private to a specific *C. briggsae* gene annotation would reflect instances of incorrectly predicted genes. The number of protein sequences with identical length (no difference from N2), near-identical length (within 5% of N2), and deviated length (greater than 5% difference from N2) serve as a heuristic to assess gene annotation accuracy. Using these estimates of protein length accuracy, we compared the CGC2 and AF16 cb4 gene annotations to the N2 gene annotations (Fig. 5, Supplementary Figure 16, Supplementary Table 12). When compared to N2 protein sequences, CGC2 had 12 more identical protein sequence length matches than AF16, whereas AF16 had 94 more near-identical protein sequence length matches than CGC2 (Supplementary Figure 16, Supplementary Table 12). Despite these small differences, the vast majority of genes with single-copy orthologs present in both *C. briggsae* gene annotations are accurate (identical or near-identical) in protein sequence length relative to N2 (CGC2: 84.1% accurate, 6,753/8,021 of single-copy orthologs; AF16 cb4: 85.2% accurate, 6,835/8,021 of single-copy orthologs) (Fig 5a-b, Supplementary Table 12). A small proportion of genes contributing towards the number of accurate N2 orthologs in CGC2 (4% of accurate genes, 273/6,753) or AF16 cb4 (5.1% of accurate genes, 355/6,835) gene annotations appear to be exclusively accurate in only one of these two annotations, highlighting a small set of mispredicted genes in both gene annotations that require further analysis and correction (Fig. 5c). Moreover, 72.4% (661/913) of genes that are deviated in protein sequence length from N2 in both *C. briggsae* annotations are concordant with each other (less than 5% difference in sequence length between AF16 and CGC2), indicative of orthologs that have generally expanded or contracted in protein sequence length between *C. elegans* and *C. briggsae*. The accuracy of the remaining protein sequences that are deviated from N2 (27.6%, 252/913) cannot be determined with this analysis of protein sequence length, because the annotations between AF16 and CGC2 are discordant (greater than 5% difference) in sequence length with each other. Overall these results indicate that the CGC2 gene annotations are comparable in accuracy and completeness to the AF16 cb4 gene annotations. Further improvement could be achieved in the near future with a targeted approach that seeks to manually examine mRNA and protein sequence alignment evidence underlying gene annotations that are present in AF16 cb4 and QX1410 but were not captured in CGC2 (414 genes), are accurate in AF16 cb4 but not CGC2 relative to N2 (355 genes), or remain deviated and discordant in protein sequence length between AF16 cb4 and CGC2 (252 genes).

**Fig. 5.**
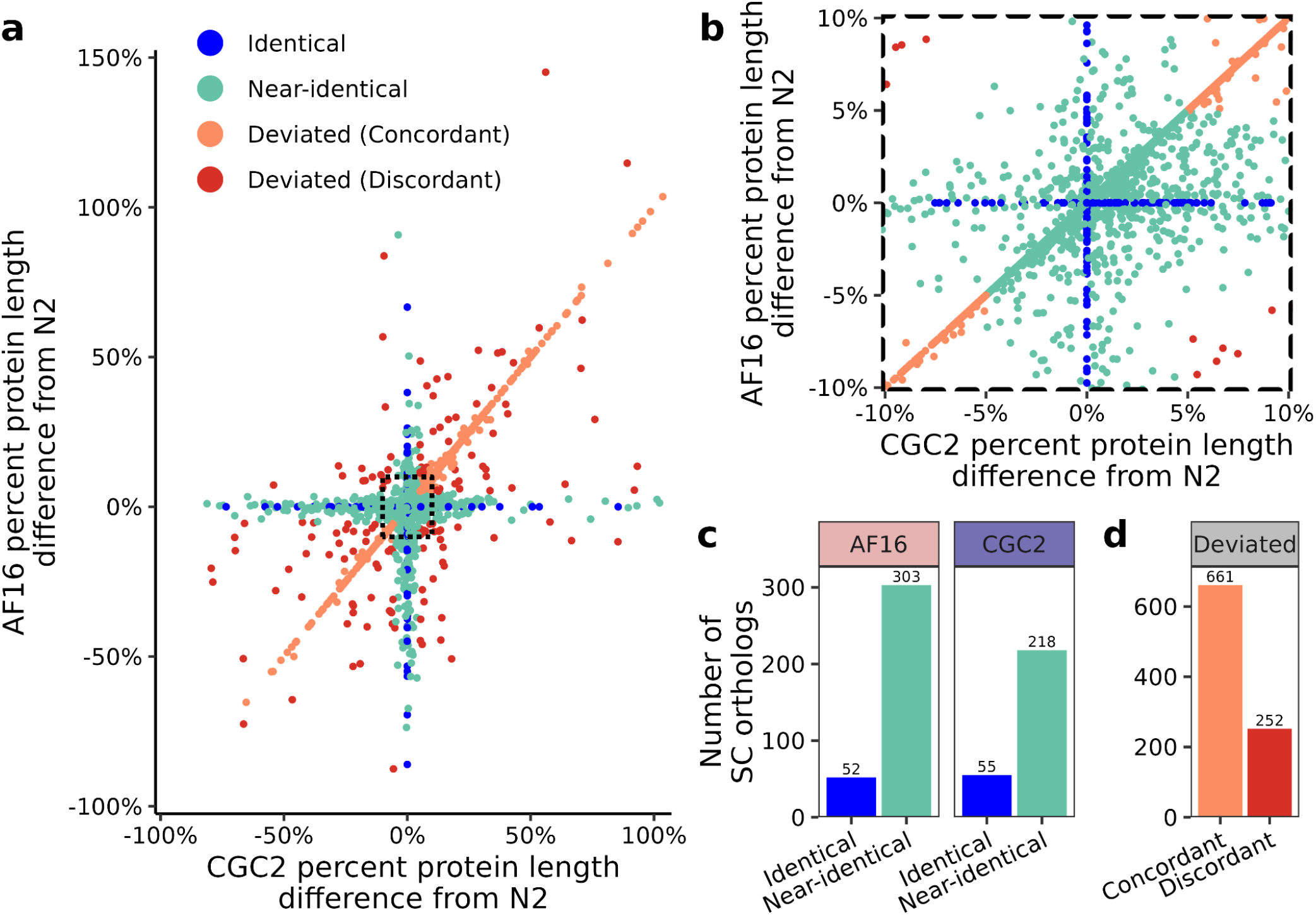
Analysis of AF16 cb4 and CGC2 gene annotations reveals high concordance in protein sequence length. (a) Percent difference between the lengths of *C. briggsae* (AF16 cb4 or CGC2) protein sequences and its orthologous *C. elegans* (N2) protein sequences. Only single-copy (SC) orthologs are included. Percent protein length difference estimates for the AF16 cb4 annotation (y-axis) relative to the CGC2 annotation (x-axis) are shown as points colored by a classification based on the length difference from N2 (identical to N2 in either *C. briggsae* annotation, blue; near-identical (less than 5% difference) to N2 in either *C. briggsae* gene annotations, teal; deviated (greater than 5% difference) from N2 but concordant between both *C. briggsae* gene annotations, orange; deviated from N2 and discordant between *C. briggsae* gene annotations, red). A total of 23 data points outside of the axes ranges were excluded for visualization clarity, but are included in the supplement (Supplementary Table 12). A black dashed square marks the range between −10% to 10% protein length difference across both axes, magnified in (b). (c) counts of genes contributing towards identical and near-identical protein length matches to N2 only in AF16 cb4 or CGC2. (d) counts of genes deviated from N2 protein sequences in both *C. briggsae* gene annotations, classified as concordant or discordant (within or greater than 5% difference) protein sequence lengths between AF16 cb4 and CGC2.

## Discussion

The AF16 reference genome built in 2003 used a combination of Sanger-based shotgun sequencing and a physical map created from fosmid sequencing and bacterial artificial chromosome libraries (Stein et al. 2003). Since that time, advances in massively parallel sequencing technologies, particularly long-read sequencing and high-throughput contact mapping technologies, in addition to optical mapping and *de novo* assembly algorithms have made reference genome creation straightforward. Efforts to update and create nematode reference genomes have been extensive, with initiatives such as the Tree of Life Programme at the Wellcome Sanger Institute and researchers who extensively curate reference genomes to create T2T assemblies. For example, since its initial sequencing in 1998, the reference genome for *C. elegans* has undergone continuous updates (The C. elegans Sequencing Consortium 1998; Hillier et al. 2005; Tyson et al. 2018; Howe 2019), with the first T2T version, CGC1, released in 2025 (Ichikawa et al. 2025). Since the initial assembly of the AF16 genome in 2003, multiple groups have released updated versions which have closed thousands of gaps and resolved megabases of previously unplaced sequence (Stein et al. 2003; Hillier et al. 2007; Ross et al. 2011; Ren et al. 2018; Xie et al. 2024; Pellow et al. 2026). In 2022, our laboratory provided a new reference genome for *C. briggsae* using an AF16-related strain, QX1410 (Stevens et al. 2022). However, despite these advances in genomic resources for *C. briggsae*, we recognized a need for an improved AF16 reference genome because of the extensive research that has been done in the AF16 genetic background over the past few decades. To create an updated AF16 reference genome assembly, we obtained high-coverage HiFi, ONT, and Hi-C sequencing data to generate a *de novo* genome assembly for an isogenic derivative of the AF16 strain, CGC2. After manual curation, CGC2 is the first gap-free, T2T genome assembly for *C. briggsae*. CGC2 is drastically more contiguous than the original AF16 assembly with captured telomeric repeats on all chromosome ends and no unplaced sequence. The CGC2 genome assembly is publicly available at GenBank and the *Caenorhabditis* Natural Diversity Resource (CaeNDR), and its source strain is available from the CGC (see Data Availability) as CGC2. In parallel to this study, a different research group recently published a gap-free, T2T assembly of AF16 (Pellow et al. 2026). Although this T2T AF16 assembly could serve as a useful resource, the history of the AF16 strain that the authors used is not supplied in their study. Additionally, the updated AF16 genome is not publicly available yet and genomic quality metrics about the updated assembly and their scaffolding and manual genome curation methods are not provided in detail. For these reasons, a comparison to CGC2 is not possible. The comparison of CGC2 to this T2T AF16 assembly will reveal potential discrepancies that arise from differences in the identity of the AF16 strains used for each assembly, in addition to *de novo* genome assembly, scaffolding, and manual curation methodologies.

To further improve the CGC2 genome assembly presented in this study, efforts are needed to resolve the rDNA cistron units on the left-end of chromosome V. This curation requires extensive manual attention and potentially ONT ultra-long sequencing to correctly orient rDNA contigs and construct the fully uncollapsed repeat sequence. Additional manual inspection of telomeric repeat sequence and fold-coverage over these repeat regions might reveal if CGC2 has completely captured the full extent of telomeric sequence. Despite these minor limitations of the CGC2 assembly, the high sequence completeness and base-calling accuracy of CGC2 in addition to its extreme contiguity will aid the design of more accurate molecular, genetic, and comparative genomics experiments.

To provide additional genomic resources for the CGC2 genome, we lifted over the genomic positions of 105 out of 108 experimentally validated indel sites and their primer sequences from the AF16 cb4 genome onto the CGC2 genome. The primer sequences of the three remaining indel variant sites mapped to multiple genomic positions so we could not effectively place them. We also exploited deep RNA-seq and protein sequence libraries to annotate protein-coding genes. Using a comparison of single-copy orthologs across *C. briggsae* and *C. elegans* reference gene annotations, we have identified 964 genes that are currently absent in the AF16 cb4 gene annotations but are present in both QX1410 and CGC2. These novel genes are not shared with *C. elegans*, reflecting *C. briggsae* specific genes that were likely underrepresented in the previous AF16 cb4 reference. We also identified 414 genes that are potentially missing in CGC2 that require additional manual validation before they can be incorporated. Lastly, although the length of translated protein sequences of both AF16 cb4 and CGC2 gene annotations are largely concordant with their *C. elegans* orthologs, we identified a small number of genes that deviate in protein sequence length from *C. elegans* in only one of the two *C. briggsae* gene annotations (AF16 cb4, 355 genes; CGC2, 273 genes). Future improvements to the CGC2 gene annotations should not only try to incorporate the missing genes from AF16 that we have identified but also examine RNA sequencing data underlying genes that are concordant in protein sequence length between AF16 and *C. elegans* gene annotations that remain deviated in CGC2. Nonetheless, we conclude that the CGC2 gene annotations are comparable in completeness and accuracy to the existing AF16 reference and will serve as a useful resource for studies of genetic variation within and across species.

*C. briggsae* now has multiple high-quality chromosome-level reference genomes, inclusive of CGC2, QX1410, and VX34, where intra- and inter-species comparative studies can be a next step. Because of the extreme levels of genetic variation within *C. briggsae* (Moya et al. 2025), comparative genomic studies among strains will elucidate synteny and conserved gene order, structural variation, and gene copy-number differences that will provide novel insights into genome and gene family evolution in genetically diverse strains. With the advent of pangenomics and high-quality reference genomes for multiple species of *Caenorhabditis* (Stein et al. 2003), inter-species comparisons of *Caenorhabditis* might reveal species-specific structural and gene copy-number variation associated with adaptation or speciation, and provide broader insights into genome evolution, such as macrosynteny conservation and the significance of programmed DNA elimination in ancestral *Caenorhabditis* species (Stevens et al. 2025).

## Data Availability

The raw DNA Illumina short-read sequencing data for PB420 is publicly available from NCBI accession PRJNA1453487. The raw PacBio HiFi data and Oxford Nanopore Technologies long-read sequencing data are publicly available from NCBI accession PRJNA1451928. The Hi-C Illumina short-read sequencing is publicly available from NCBI accession PRJNA1453590. Short-read RNA sequencing data is publicly available from NCBI accession PRJNA1454097. The final CGC2 assembly is publicly available on NCBI under accession PRJNA1451919 and the CaeNDR. The CGC2 gene annotation files are publicly available on CaeNDR. Scripts and related raw and processed data are available at GitHub (https://github.com/AndersenLab/CGC2_genome_MS)

## Acknowledgements

We thank Drs. Scott Baird, Helen Chamberlin, and Bhagwati Gupta for AF16 derivative strains. We thank the *Caenorhab*ditis Genetics Center, which is funded by the National Institute of Health Office of Research Infrastructure Programs (P40 OD010440). We thank WormBase, which is supported by grant #U24 HG002223 from the National Human Genome Research Institute, the UK Medical Research Council and the UK Biotechnology and Biological Sciences Research Council. We thank the *Caenorhabditis* Natural Diversity Resource, which was supported by the National Science Foundation Capacity grant (2224885). We thank members of the Andersen lab for their feedback on the manuscript.

## Author Contributions

L.M.O., N.D.M., N.S.J., and E.C.A. conceived and designed the study.

L.M.O. and N.D.M. analyzed the data.

L.M.O., N.D.M., N.S.J., and E.C.A. wrote the manuscript.

R.E.T., S.E.B., and H.M.C. edited the manuscript.

R.E.T., A.K., and N.S.J. performed whole-genome sequencing for CGC2.

H.M.C. and S.E.B. provided AF16 derivative strains

H.M.C. and H.L. performed indel validations

