## Supplementary Figures for "A gap-free, telomere-to-telomere genome assembly for the *Caenorhabditis briggsae* reference strain AF16"

**A gap-free, telomere-to-telomere genome and improved gene models for the *Caenorhabditis briggsae* reference strain AF16**

Lance M. O'Connor[1,2], Nicolas D. Moya[1], Nikita S. Jhaveri[1], Robyn E. Tanny[1], Ayeh Khorshidian[1], Haimeng Lyu[3], Helen M. Chamberlin[3], Scott E. Baird[4], and Erik C. Andersen[1]*

### SUPPLEMENTARY FIGURES

**Supplementary Figure 1:** Alternative alleles across AF16 derivatives.
**Supplementary Figure 2:** Whole-genome alignment of AF16 cb5 to QX1410.
**Supplementary Figure 3:** Blob plots of CGC2 contig taxonomy classification.
**Supplementary Figure 4:** Identification of the duplicate haplotig, scaffold 7.
**Supplementary Figure 5:** Sequencing reads alignment to CGC2 chromosome III gap in IGV.
**Supplementary Figure 6:** Hi-C contact map of CGC2 scaffolds.
**Supplementary Figure 7:** Genome-genome alignments of CGC2 to QX1410 highlighting scaffold gaps.
**Supplementary Figure 8:** Nuclear chromosome sizes of AF16 and CGC2.
**Supplementary Figure 9:** A shared inversion in VX34 and AF16 reference genomes on the arm of QX1410 chromosome V.
**Supplementary Figure 10:** A private QX1410 inversion allele.
**Supplementary Figure 11:** Telomeric repeat sequence in *C. briggsae* genome assemblies.
**Supplementary Figure 12:** Long-read sequencing coverage on the left-end of CGC2 chromosome V.
**Supplementary Figure 13:** Whole-genome alignment of AF16 (cb4) to CGC2.
**Supplementary Figure 14:** Multi-mapping insertion-deletion primers.
**Supplementary Figure 15:** Inter-chromosomal single-copy ortholog mappings between *C. briggsae* genomes.
**Supplementary Figure 16:** Comparison of protein length accuracy of *C. briggsae* single-copy orthologs

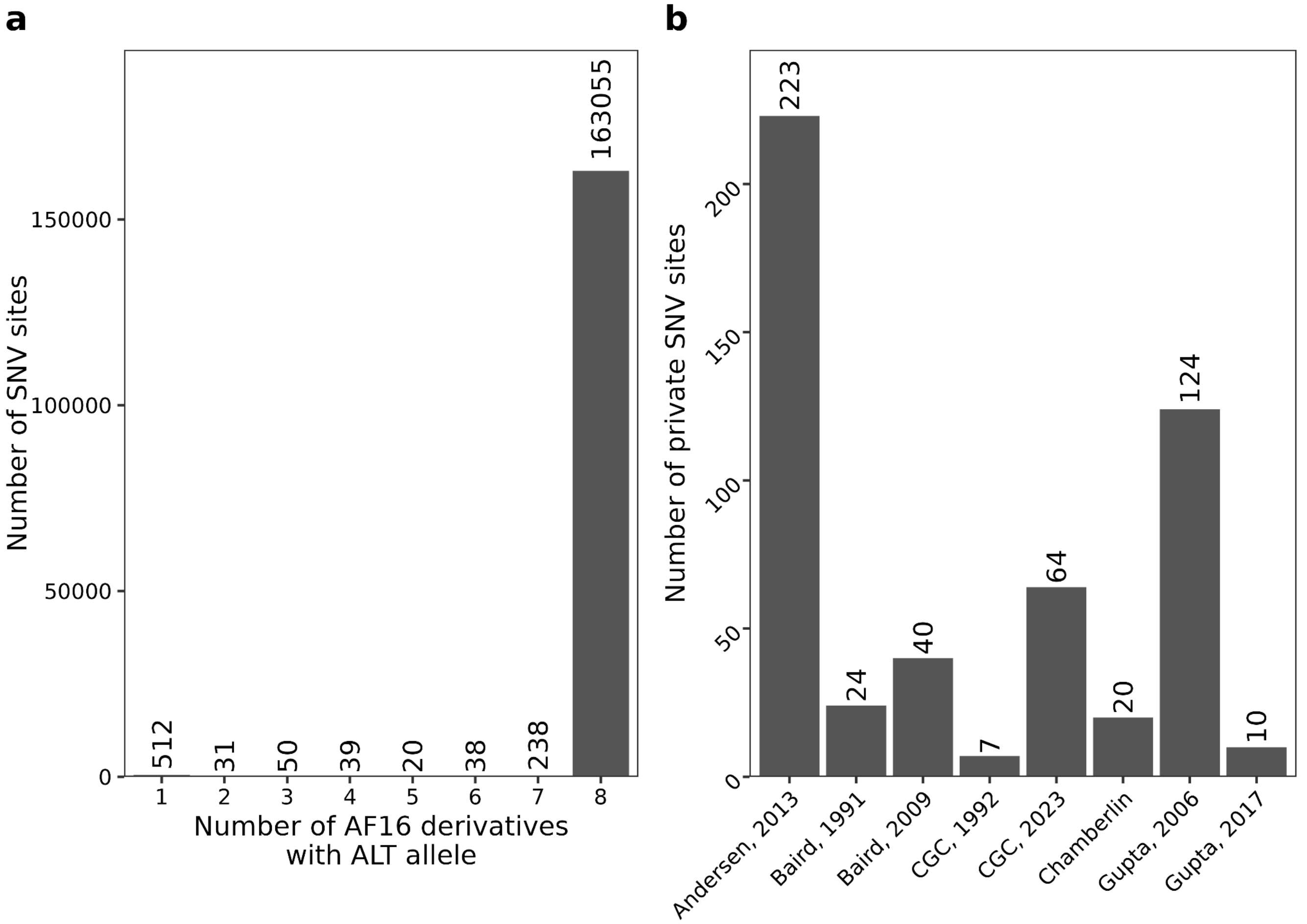


**Supplementary Figure 1.** Alternative alleles across AF16 derivatives. Only SNV sites with perfect information (no missing genotypes) are considered. (a) Bar chart showing the number of SNV sites where one or more AF16 derivative strains carry an alternative (ALT) allele relative to the QX1410 reference genome. All eight AF16 derivatives carry an ALT allele at the vast majority (163,055) of SNV sites, representing natural variants between the AF16 derivatives and QX1410. A total of 512 sites are private to only one AF16 derivative. (b) Bar chart showing the number of ALT alleles at private SNV sites across AF16 derivative strains (labels display source laboratory and the year that the strain was cryopreserved.

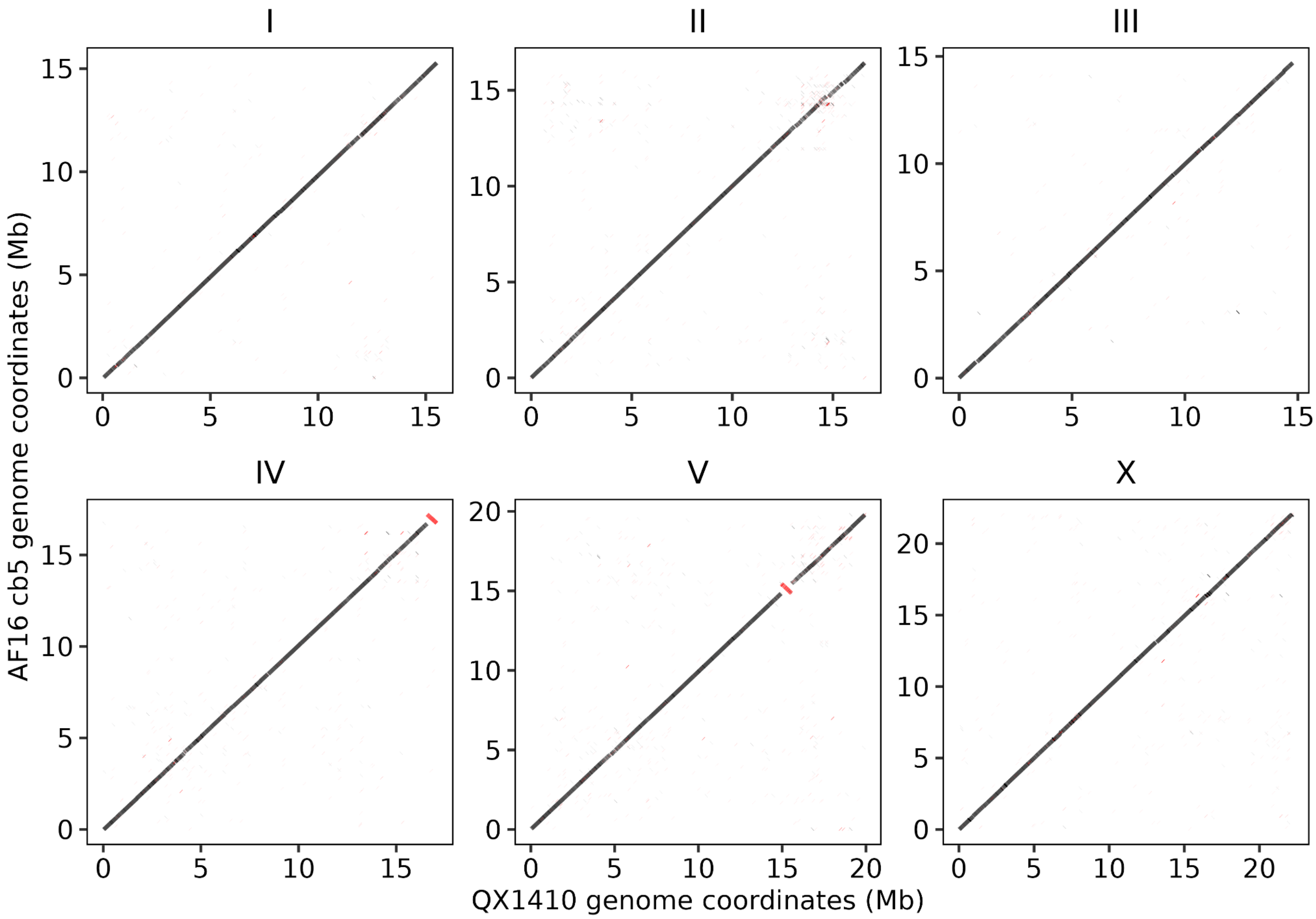


**Supplementary Figure 2.** Whole-genome alignment of AF16 cb5 to QX1410. Colinearity of AF16 cb5 to QX1410 genomes generated using nucmer. Inter-chromosomal alignments and alignments spanning less than 1 kb are not shown. Alignments in the reverse orientation are displayed in red.

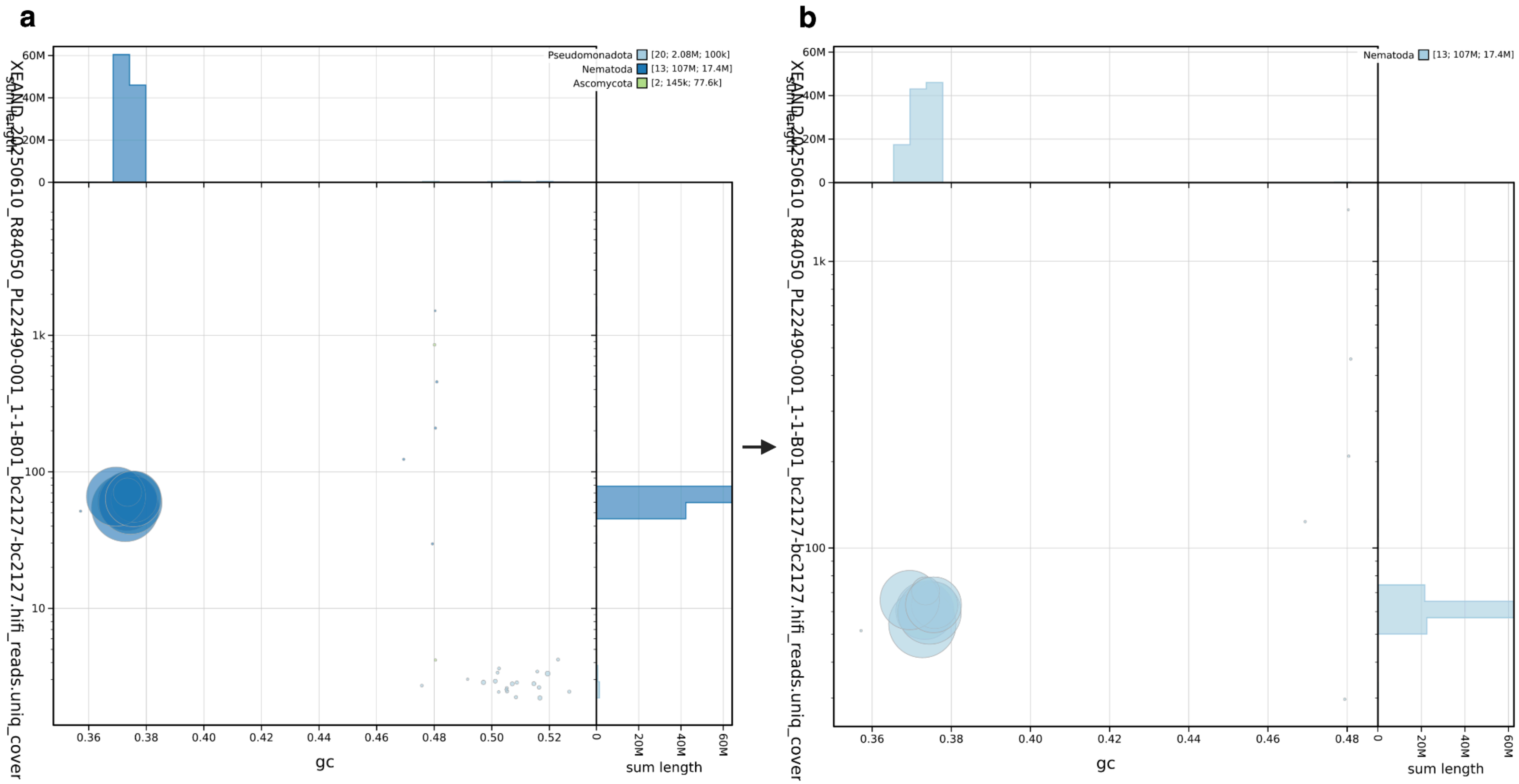


**Supplementary Figure 3.** Blob plots of CGC2 contig taxonomy classification. (a) A blob plot produced using BlobToolKit. Contigs are colored by their taxonomic classification. The x-axis displays contig GC content, and the y-axis displays HiFi read coverage of contigs. (b) The resulting blob plot of 13 contigs after removing contigs that were not classified as *Nematoda*.

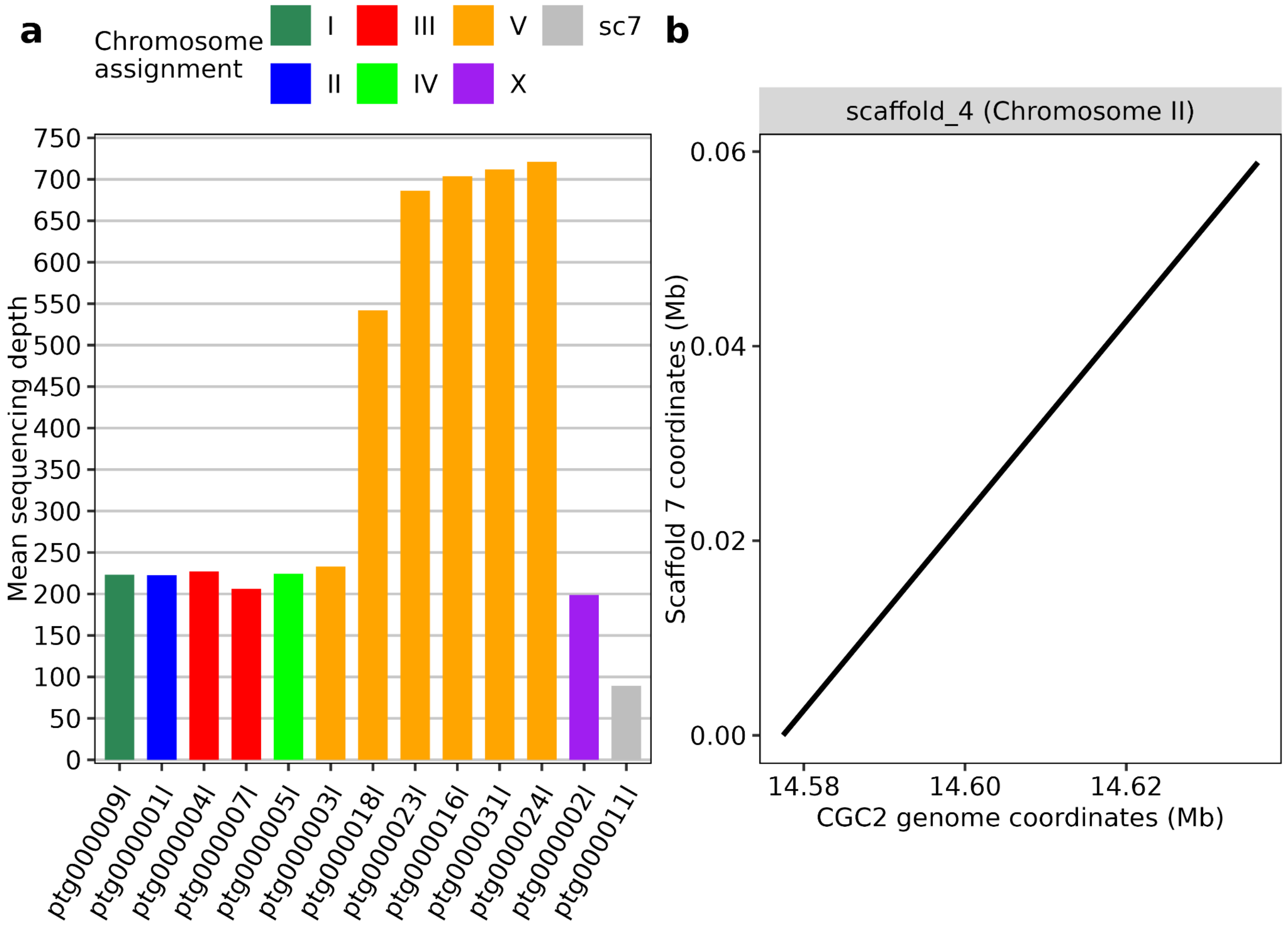


**Supplementary Figure 4.** Identification of the duplicate haplotig, scaffold 7. (a) Hi-C read fold-coverage of the 13 contigs used in scaffolding. Contigs are colored by their final chromosome identification after scaffolding, with scaffold 7 (sc7) corresponding to a single contig. (b) Alignment of scaffold 7 to CGC2 scaffolds, where scaffold 7 aligns to a region in scaffold 4 (chromosome II).

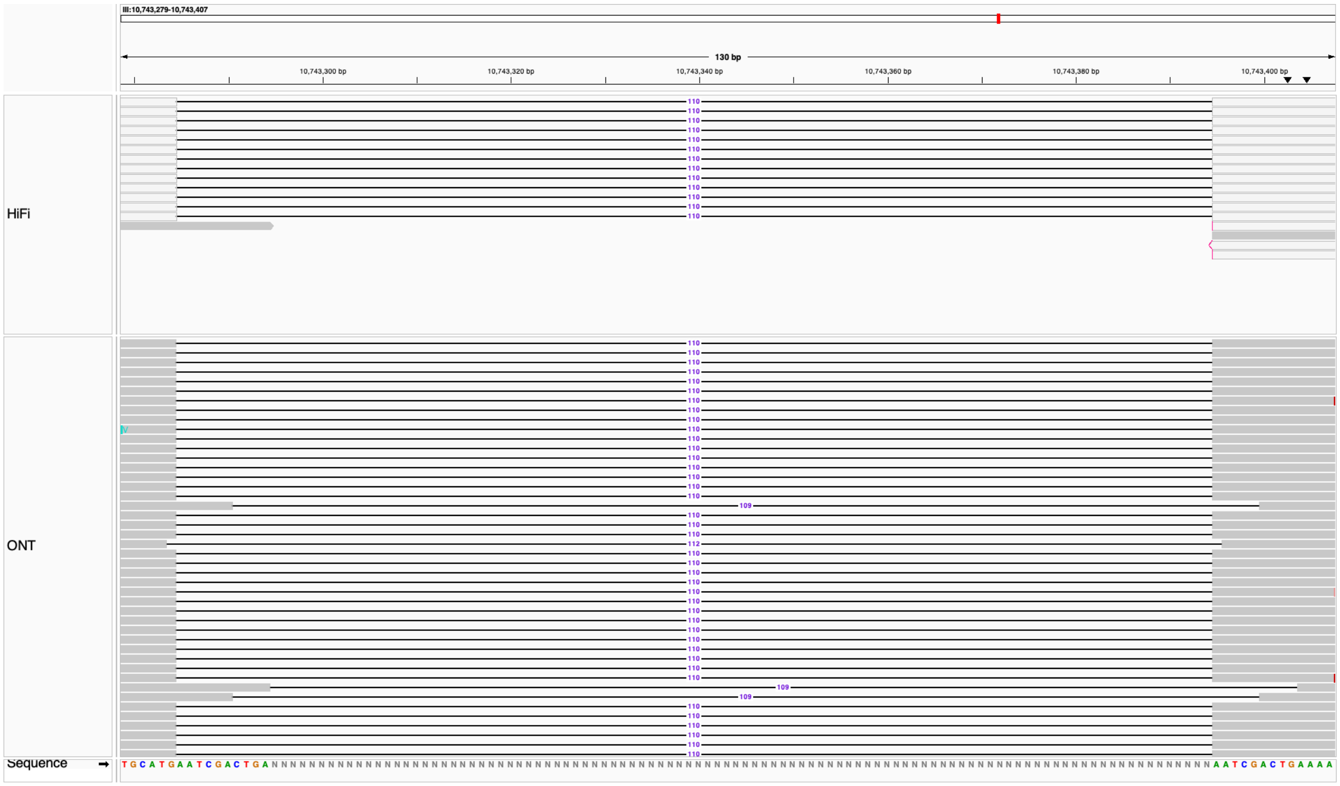


**Supplementary Figure 5.** Long-read sequencing reads alignments to CGC2 chromosome III gap in IGV. The first row displays PacBio HiFi read alignments and the second row displays ONT read alignments. The gap is represented with 100 Ns, displayed in the bottom sequence panel, where the last 100 bps upstream of the gap and the first 100 bps downstream of the gap are identical. The removal of the 100 Ns and the 10 bps upstream of the gap is supported by multi-mapping HiFi reads and MAPQ60 ONT reads. ONT reads are sorted in descending order of mapping quality and grey-filled reads indicate primary read alignments.

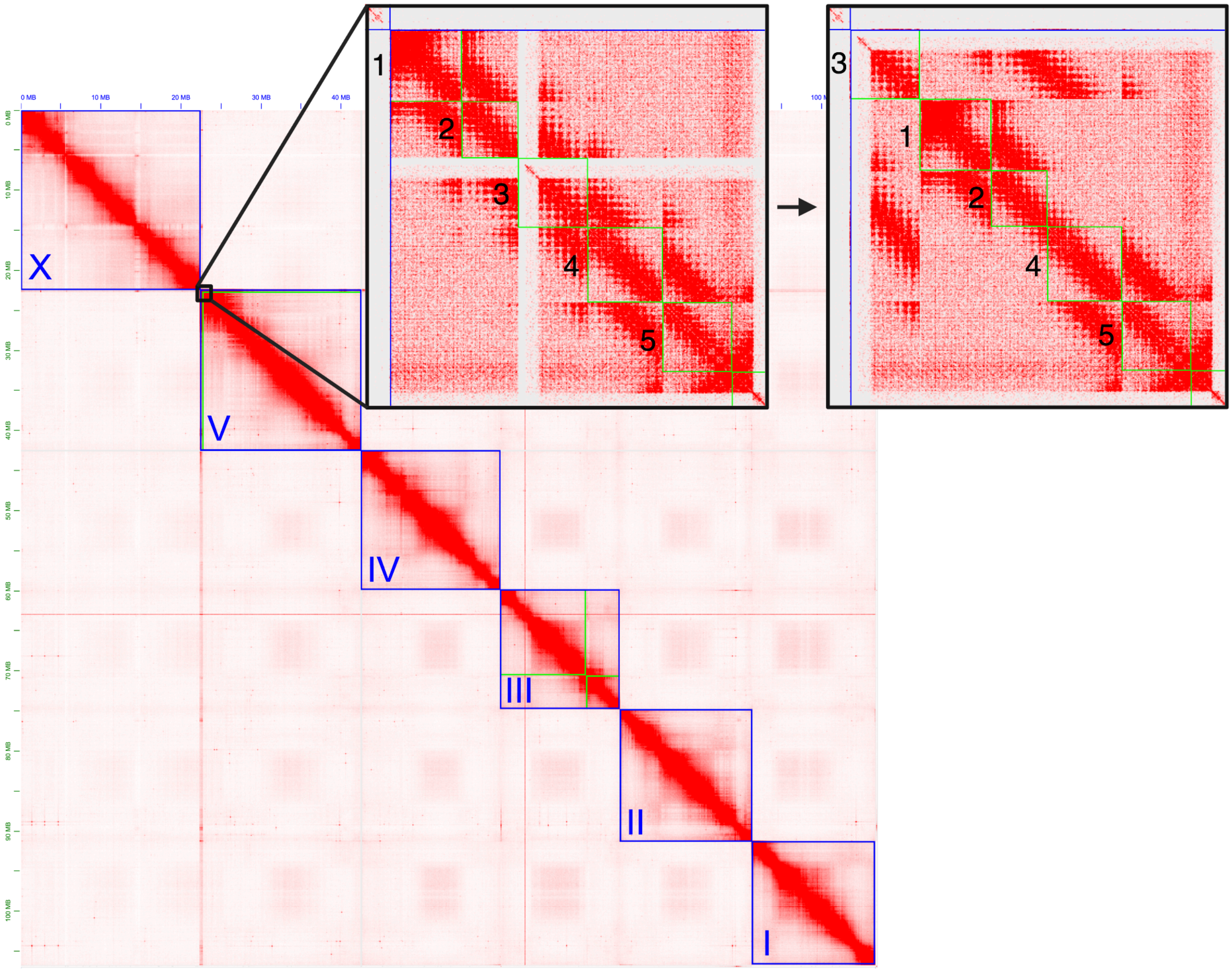


**Supplementary Figure 6.** Hi-C contact map of CGC2 genome assembly. The diagonals inside blue squares represent nuclear chromosomes and the diagonals inside smaller, green squares represent contigs. Chromosome III is constructed from two contigs, where chromosome V is constructed from six contigs. The left inset displays five labeled contigs at the left-end of chromosome V, where five gaps are present (vertices of green squares). The right inset displays the resulting contact map after moving contig three, which contains telomeric repeat sequence, to the terminal left-end of chromosome V.

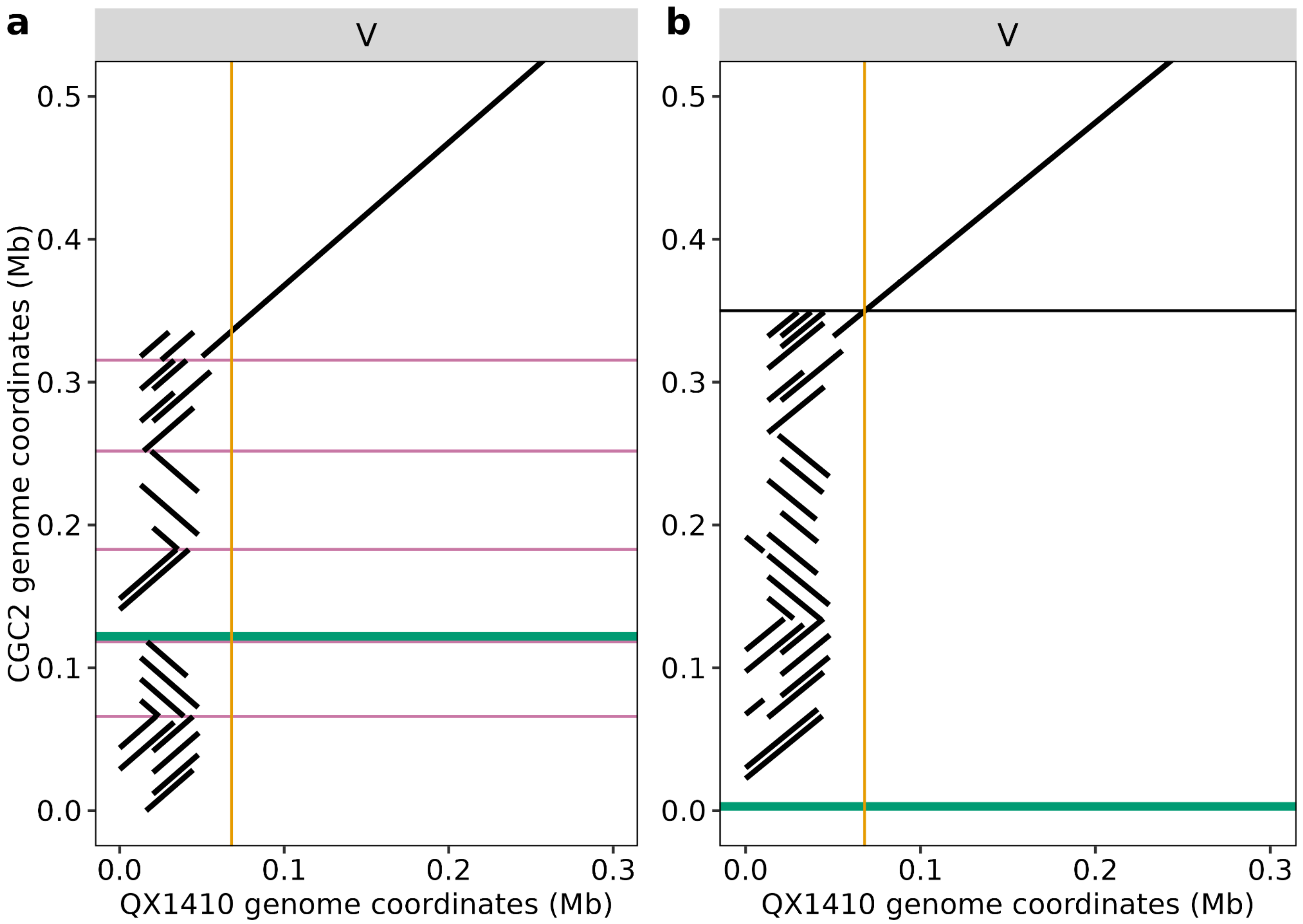


**Supplementary Figure 7.** Genome-genome alignments of CGC2 to QX1410 highlighting scaffold gaps. (a) Left-end of chromosome V displaying the remaining gaps (horizontal purple lines), telomeric repeat sequence (horizontal green line), and the terminus of rDNA cistron repeats in QX1410 (orange vertical line). (b) The same plot as panel (a) after contig-reordering to place telomeric repeat sequence at the left-end of chromosome V based on Hi-C contact evidence and gap closing using TGS-GapCloser (Xu et al. 2020).

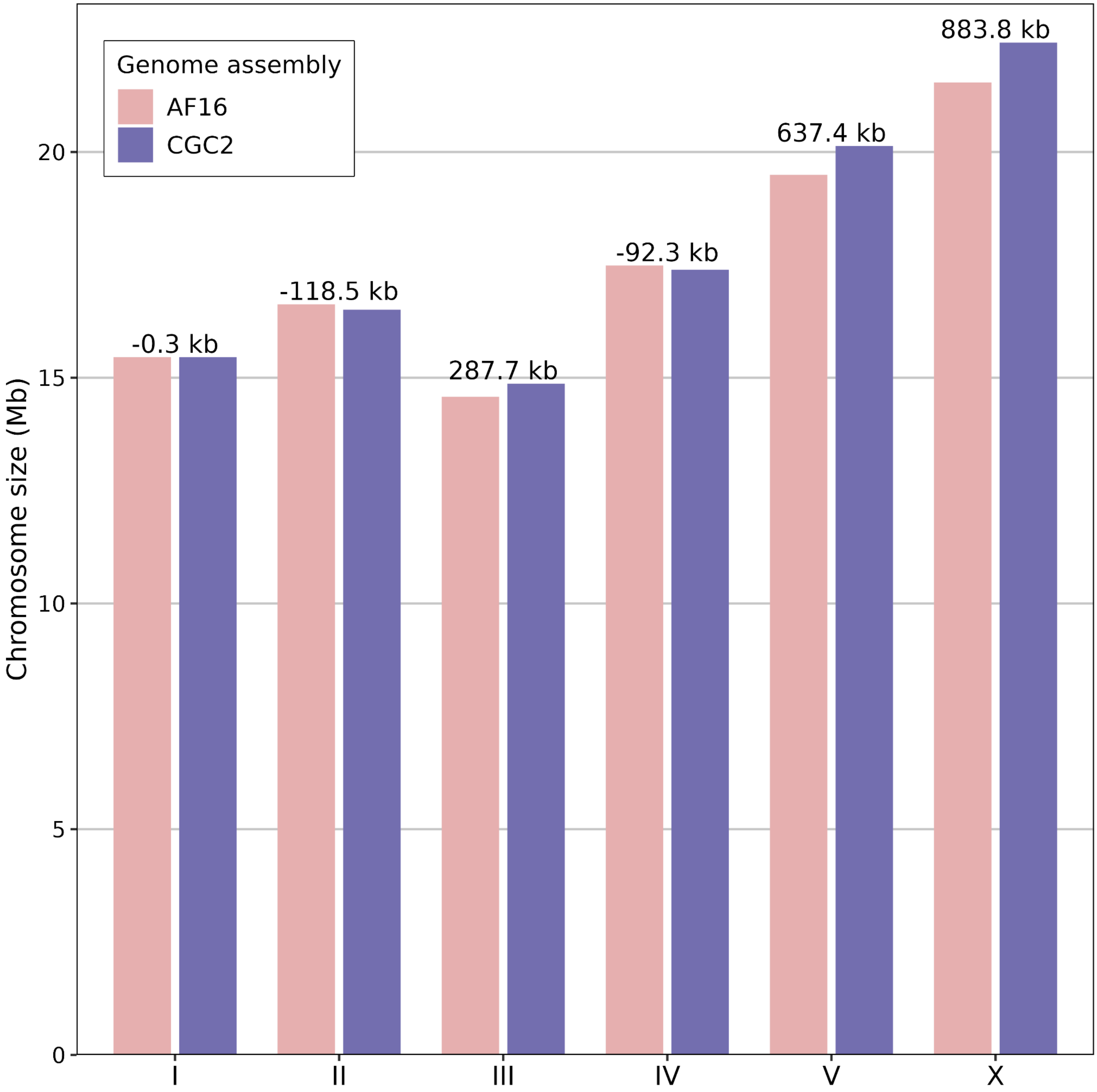


**Supplementary Figure 8.** Nuclear chromosome sizes of AF16 cb4 and CGC2. Differences in nuclear chromosome sizes of CGC2 compared to AF16 cb4 are displayed above each chromosome set, where the largest difference in span is chromosome X, and smallest difference in span is chromosome I.

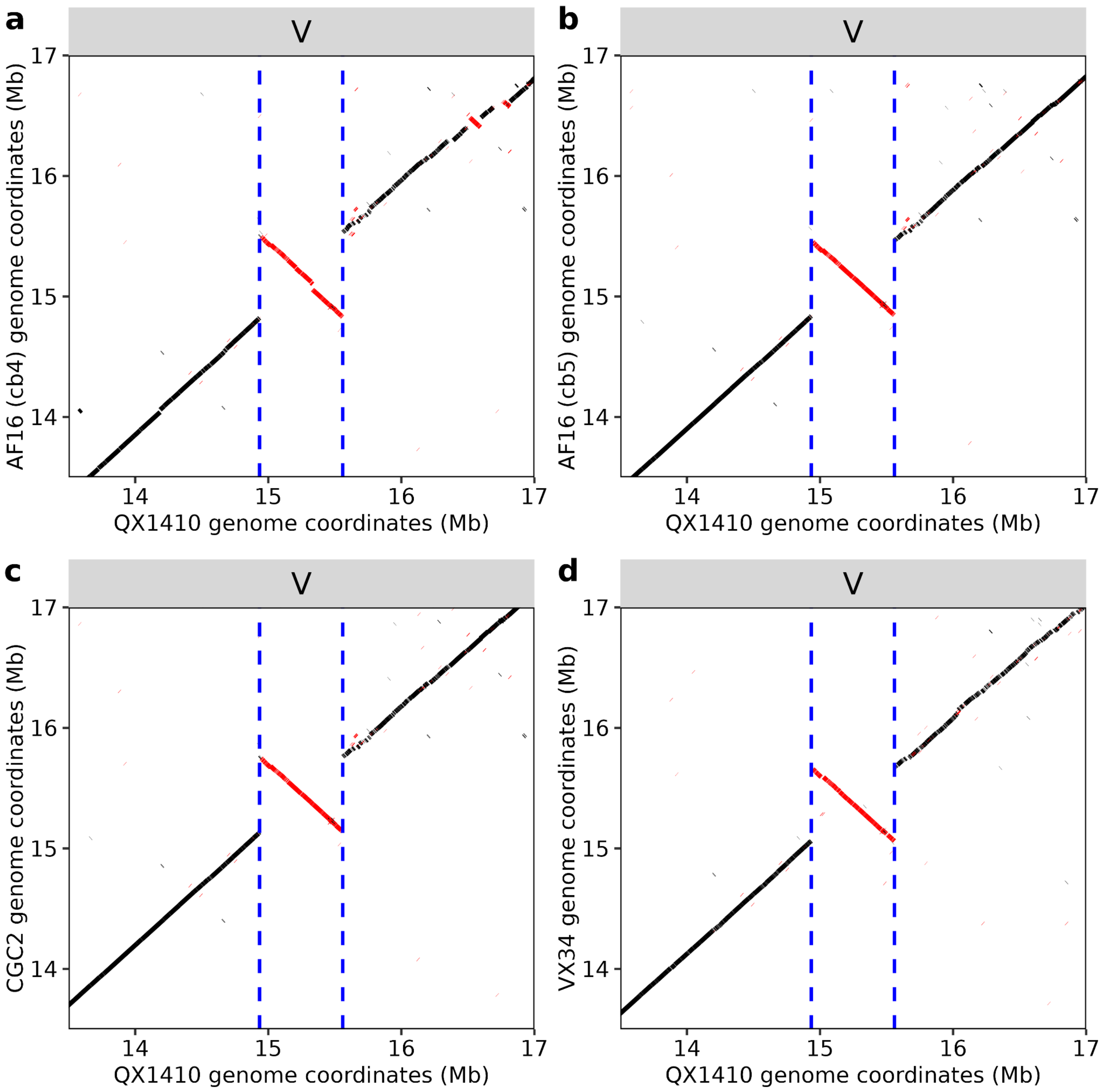


**Supplementary Figure 9.** A shared inversion in VX34 and AF16 reference genomes on the arm of QX1410 chromosome V. (a) AF16 (cb4) genome alignments to QX1410 highlighting an inversion (red) on the right arm of chromosome V. Vertical blue dashed lines indicate the breakpoints of the inversion in QX1410 genome coordinates. Genome-genome alignments of AF16 (cb5) (b), CGC2 (c), and VX34 (d) to QX1410 highlighting the same inversion (red) found in AF16 (cb4). Vertical blue dashed lines are the same coordinates in all panels.

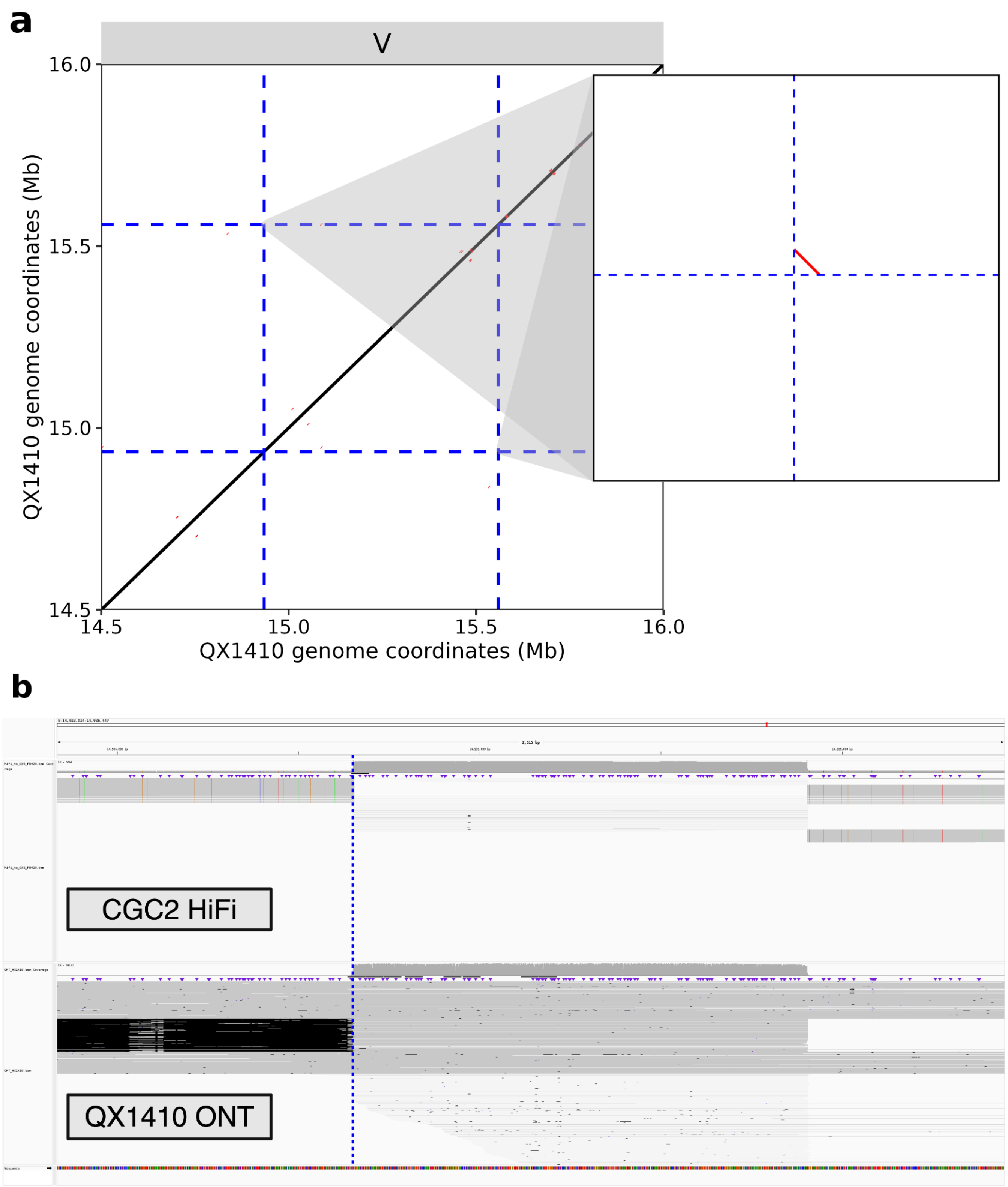


**Supplementary Figure 10.** A private QX1410 inversion allele. a) Whole-genome alignments of QX1410 to itself at the locus that is inverted in other *C. briggsae* reference genomes (Supplementary Figure 9). Inset displays multi-aligning sequence flanking the inversion breakpoints (dashed blue lines) b) IGV screenshot of CGC2 PacBio HiFi read alignments (first row) and QX1410 ONT read alignments (second row) to the QX1410 genome over the left inversion breakpoint in QX1410 (vertical dashed blue line). High-quality read mappings are shaded in grey.

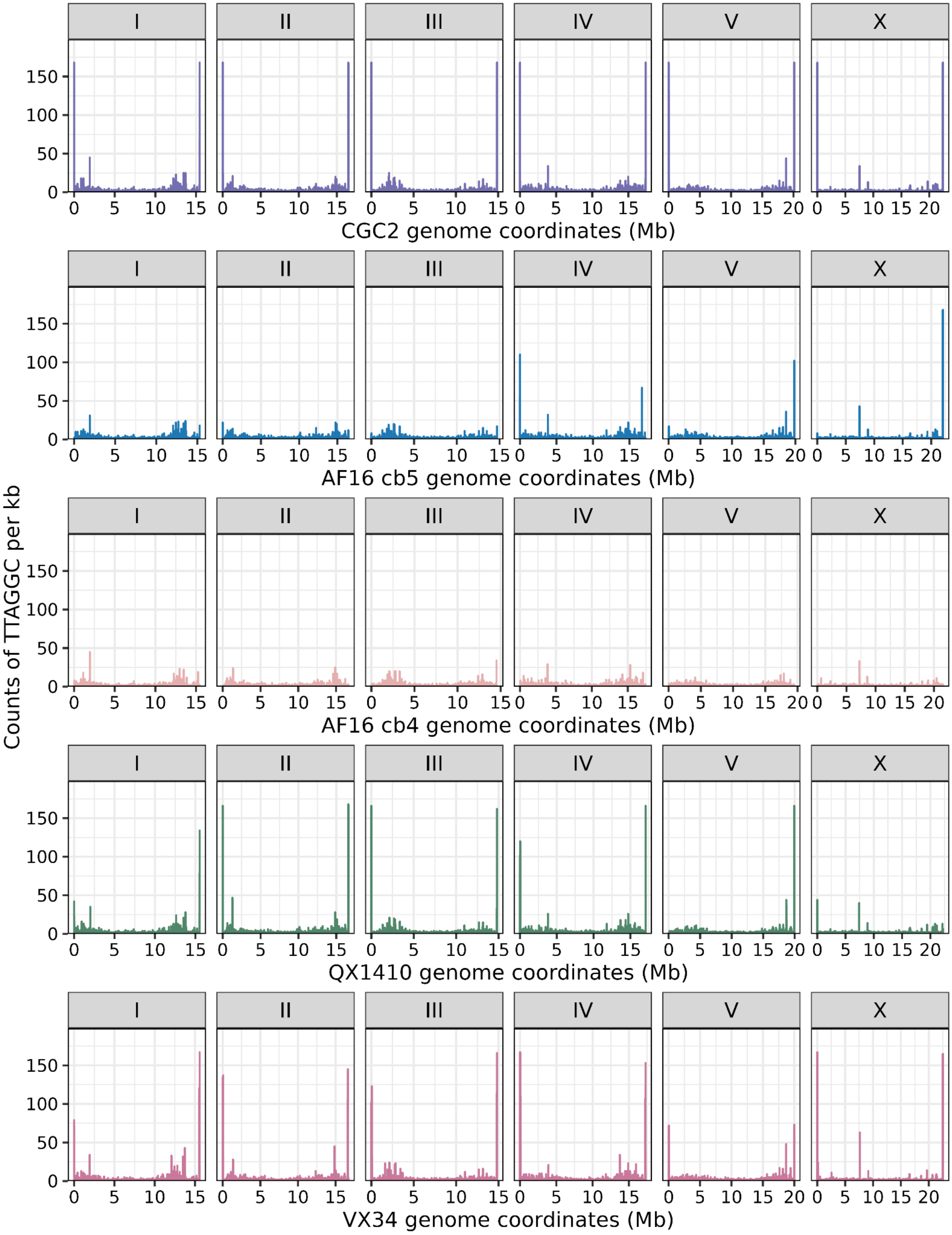


**Supplementary Figure 11.** Telomeric repeat sequence in *C. briggsae* genome assemblies. Counts of nematode telomere repeat sequence (TTAGGC) per kb bin in *C. briggsae* chromosome-level genome assemblies. Counts of 167 represent telomeric repeat sequence spanning an entire kb window.

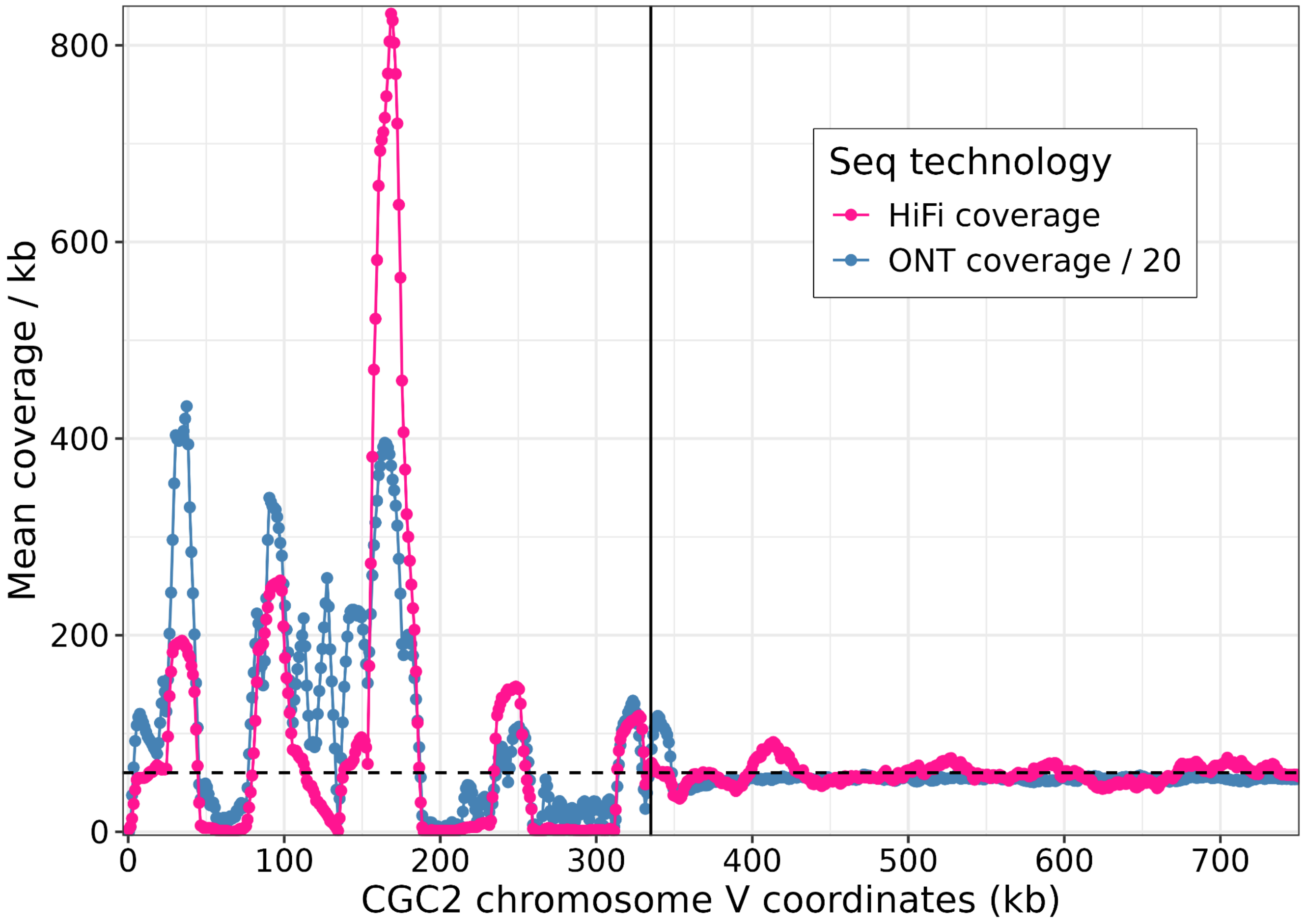


**Supplementary Figure 12.** Long-read sequencing coverage on the left-end of CGC2 chromosome V. HiFi and ONT read coverage over telomeric and rDNA repeat regions (0 kb - 335 kb (vertical black line)).

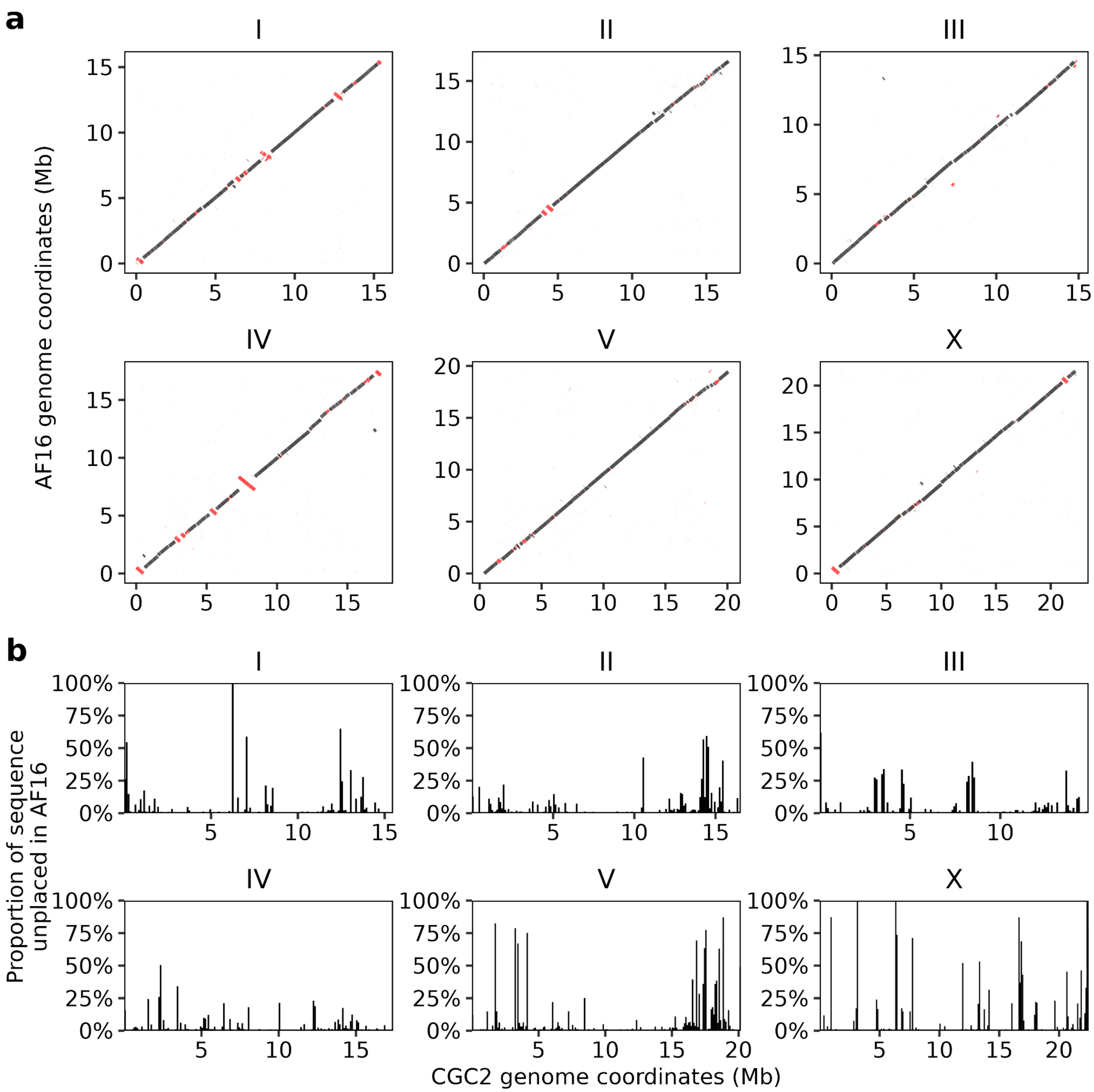


**Supplementary Figure 13.** Whole-genome alignment of AF16 cb4 to CGC2. (a) Colinearity of AF16 cb4 to CGC2 genomes generated using nucmer. Inter-chromosomal alignments and alignments spanning less than 1 kb are not shown. Alignments in the reverse orientation are displayed in red. (b) The proportion of sequence that is contained in AF16 cb4 unplaced scaffolds per 100 kb window across the CGC2 genome.

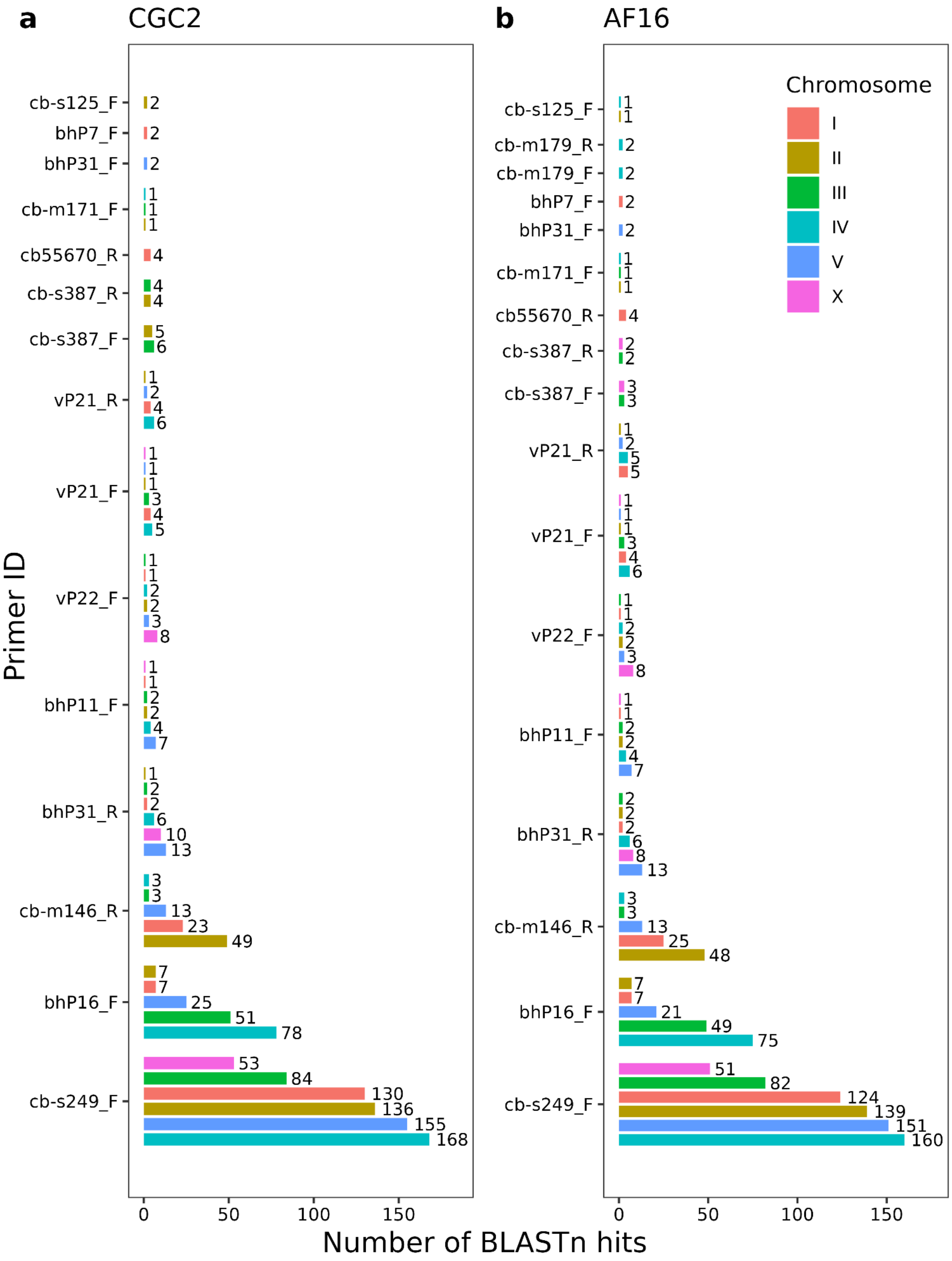


**Supplementary Figure 14.** Multi-mapping insertion-deletion primers. Bar charts showing the number of BLASTn hits (x-axis) for each primer (y-axis) with more than two mapping locations in the (a) CGC2 genome or (b) AF16 cb4 genome. Primers with multiple bars indicate that it was mapped to multiple chromosomes, with bars colored by chromosome ID.

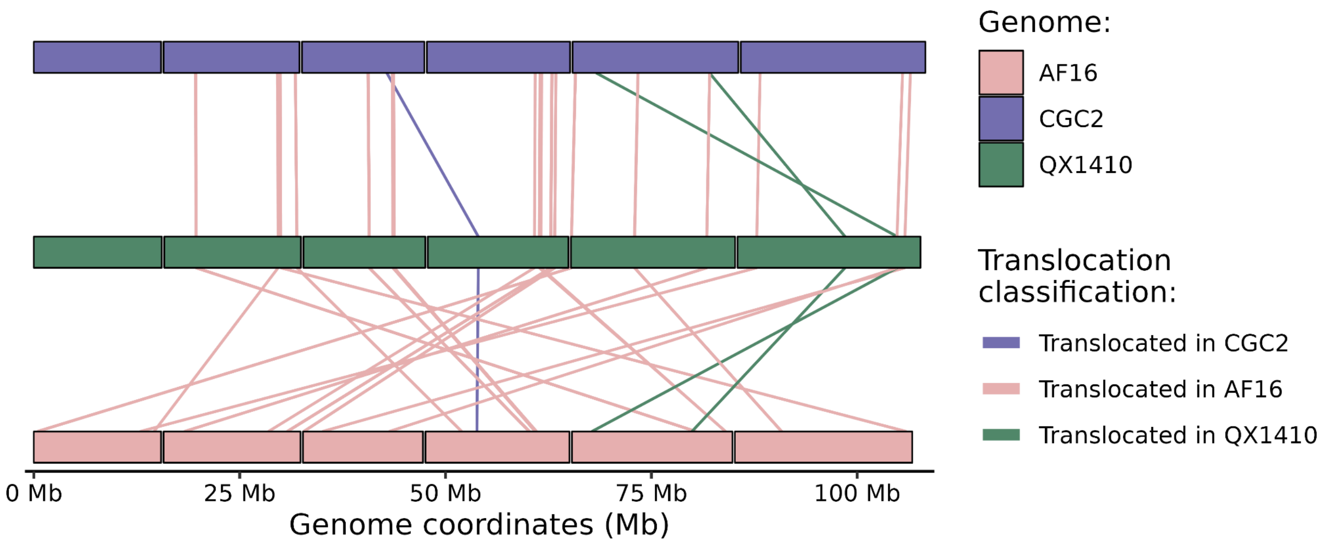


**Supplementary Figure 15.** Inter-chromosomal single-copy ortholog mappings between *C. briggsae* genomes. Each bar represents a *C. briggsae* chromosome colored and organized in rows by reference genome (CGC2, purple; QX1410, green; AF16 cb4, pink). Genome position in megabases is displayed on the x-axis. Lines represent the positions of single-copy ortholog mappings across any two genomes. Lines colored in pink represent single-copy orthologs that map within the same chromosome in CGC2 and QX1410 but map to a different chromosome in AF1 cb46. Lines colored in green represent single-copy orthologs that map within the same chromosome in CGC2 and AF16 cb4 but map across to a different chromosome in QX1410. Lines colored in purple represent single-copy orthologs that map within the same chromosome in QX1410 and AF16 cb4 but map to a different chromosome in CGC2.

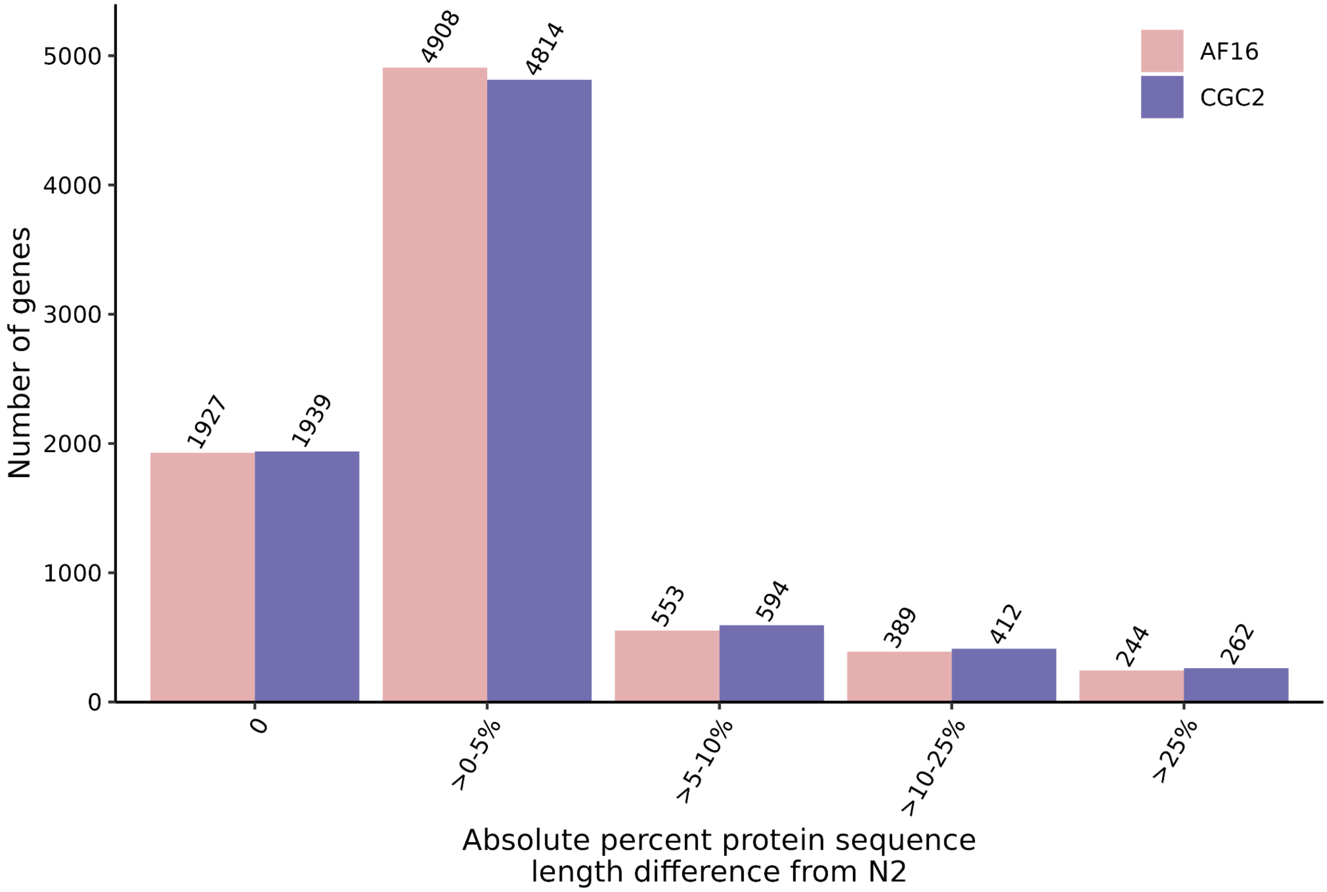


**Supplementary Figure 16.** Comparison of protein length accuracy of *C. briggsae* single-copy orthologs. Bar charts showing the number of single-copy orthologs between *C. briggsae* gene annotations (AF16 cb4, pink; CGC2, purple) across different bins of percent protein sequence length difference from single-copy *C. elegans* N2 orthologs. Identical protein sequences between *C. briggsae* and *C. elegans* will have a percent difference of zero, increasing in value depending on how much shorter or longer the *C. briggsae* protein sequence is from its N2 ortholog. Only single-copy orthologs present across all three AF16 cb4, CGC2, and N2 are included.
